# Moderate-term dimethyl fumarate treatment reduces pathology of dystrophic skeletal and cardiac muscle in a mouse model

**DOI:** 10.1101/2024.07.13.601627

**Authors:** Stephanie Kourakis, Cara A. Timpani, Ryan M. Bagaric, Bo Qi, Benazir A. Ali, Rebecca Boyer, Guinevere Spiesberger, Nitika Kandhari, Amanda L. Peterson, Didier Debrincat, Thomas J. Yates, Xu Yan, Jujiao Kuang, Judy B. de Haan, Nicole Stupka, Brunda Nijagal, Deanna Deveson-Lucas, Dirk Fischer, Emma Rybalka

## Abstract

In Duchenne muscular dystrophy (DMD), corticosteroids significantly slow disease progression and have been used as a standard of care tool for more than 30 years. However, corticosteroids also impart side effects severe enough to preclude use in some patients. There remains an unmet need for new therapeutics that target the flow-on pathogenic mechanisms of DMD with a more favourable side-effect profile. We have previously demonstrated that short-term treatment with dual-purpose anti-inflammatory, anti-oxidative dimethyl fumarate (DMF), a drug with indication and established safety data in Multiple Sclerosis, more selectively modulates Duchenne (*mdx*) immunology than the frequently used corticosteroid, prednisone (PRED). Here, we assess the effect of moderate-term DMF treatment over 5 weeks in the typically mild *mdx* mouse model that we aggravated using exercise. We show that like PRED, DMF maintains anti-inflammatory action but with additional anti-fibrotic and anti-lipogenic effects on muscle with moderate-term use. This study supports our previous work highlighting DMF as a possible repurposing candidate for DMD, especially for patients who cannot tolerate chronic corticosteroid treatment.

## Introduction

Duchenne muscular dystrophy (DMD) is a severe, progressive, muscle-wasting disease caused by mutations in the *DMD* gene. Dystrophin loss impairs the junction between the intracellular cytoskeleton and extracellular matrix (ECM), which stabilises the muscle fibre during contraction. Lack of dystrophin results in chronic muscle damage, degradation, fibrosis and replacement of skeletal muscle with fatty infiltrates (1). Affecting predominantly males, DMD is typically diagnosed by ~5 years of age presenting as delayed motor milestones and frequent falls (2) and by early adolescence, loss of ambulation and pulmonary function decline is common (3–5). Pulmonary distress is complicated by diaphragmatic weakness and fibrosis, and scoliosis, which alters the shape of the thoracic cavity and reduces respiratory capacity (6). Natural history data suggest that cardiac symptoms can emerge from as early as 12-14 years leading to dilated cardiomyopathy in almost all patients even with prophylactic treatment with cardiac medication, which can delay the onset.

The DMD therapeutic landscape has grown significantly over the past decade. Since 2016, four exon-skipping antisense oligonucleotide drugs (Exondys 51, Vyondys 53, Viltepso and Amonndys 45) and two gene replacement therapies (Elevidys and fordadistrogene movaparvovec) have been granted accelerated access (7). However, their efficacy is unclear, and fatalities have occurred with gene replacement strategies raising safety concerns (8, 9). While these therapies may have the best potential of providing long-term disease-modifying benefits, there are relatively fewer therapeutics in development to address the secondary pathomechanisms, especially the hyper-inflammation and immune system activation that drives muscle fibrosis (10). Anti-inflammatory corticosteroids have prevailed as the pharmacological standard of care (SOC) for DMD for over three decades targeting this aspect. With early intervention, they effectively slow disease progression, prolonging ambulation, and pulmonary function yet they have an extensive side-effect profile, e.g., weight gain, osteoporosis and metabolic syndrome (11). Approved in 2023, steroid analogue Agamree (vamorolone) appears to reduce some of these side-effects. However, the typical corticosteroid side effects of weight gain, behavioural issues, adrenal suppression, and cushingoid symptoms persist (12, 13), indicating that non-steroid interventions may be better for long-term use. Recently approved non-steroidal anti-inflammatory, Duvyzat (givinostat), a pan-histone deacetylase (HDAC) inhibitor was showed to slow loss of muscle strength (14) yet significant side-effects were reported in the Phase III trial (e.g. diarrhoea, vomiting, thrombocytopenia) indicating a well-balanced efficacy-to-safety profile is critical.

There has been interest in therapeutics that target molecular mechanisms outside of the genetic component in DMD including disrupted cellular homeostasis and subsequent mitochondrial dysfunction, chronic inflammation, and the replacement of muscle tissue with fibrosis and non-functional adipose tissue (15). We recently identified dimethyl fumarate (DMF), a drug with established safety data, prescribed for the autoimmune diseases, relapsing-remitting multiple sclerosis (MS) and psoriasis (15), as a repurposing candidate that can target these aspects in DMD owing to its unique mechanism of action (MOA). DMF is a potent immunoregulatory, anti-inflammatory and -oxidative drug via nuclear factor erythroid 2-related factor 2 (Nrf2) activation and hydroxycarboxylic acid receptor 2 (HCAR2) agonism (15). Our proof-of-concept study in the *mdx* mouse demonstrated short-term DMF treatment was more effective against key disease indices than SOC prednisone (PRED), including histopathology, muscle function, mitochondrial metabolism and disease driving gene signatures (16).

The *mdx* mouse is a useful pre-clinical therapeutic screening tool but presents with challenges that may compromise translational outcomes later in the drug development pipeline. After an initial acute degenerative period during rapid maturational growth at ~3-5 weeks age, the disease stabilises into relatively mild cyclical degeneration and regeneration bouts across the lifespan until ~12 months age when disease progression escalates (17). Applied exercise exacerbates muscle dystrophy in the *mdx* mouse (18), particularly fibrosis and subsequent lipid infiltration. Here, we used moderate exercise to aggravate disease and better recapitulate the DMD phenotype through which to scope DMF’s efficacy and translational potential. We aimed to make a head-to-head comparison between DMF and SOC PRED as well as assess potential additive effects through combinatorial treatment.

## Results

### Exercise aggravation increases plasma creatine kinase (CK) levels and worsens dystrophic muscle histopathology

The dystrophic *mdx* mouse is the most used pre-clinical drug screening tool despite its relatively mild phenotype (Figure 1A) compared to patients. Phenotype differences between sedentary wild-type (WT) and *mdx* mice are obvious at 8 weeks age, including elevated plasma (Figure 1B) and skeletal (Figure 1D-H, J-Q) and cardiac (Figure 1I & R) muscle damage indicators. Implementation of TREAT-NMD’s standard operating procedures enables the standardisation of pre-clinical data across laboratory groups for improved translational outcomes. We used twice-weekly forced horizontal treadmill running from the age of 35 days to aggravate the *mdx* phenotype (Figure 1A) according to TREAT-NMD SOP DMD_M.2.1.001 (19).

**Figure 1.**
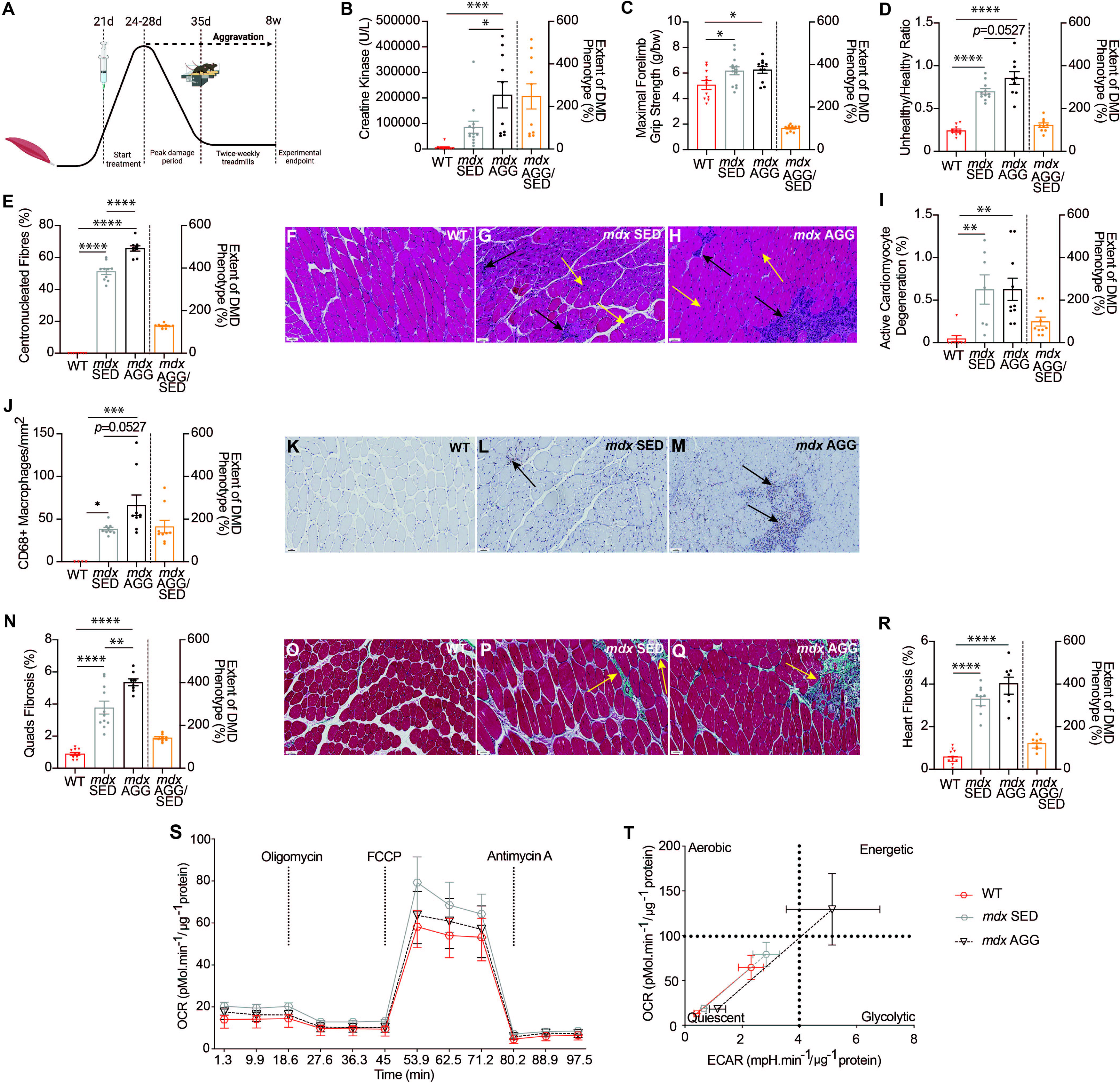
Twice-weekly forced treadmill running aggravates the *mdx* phenotype. **(A)** Schematic of treatment and treadmill protocols. *Mdx* aggravation was proved using clinically compatible parameters of **(B)** plasma creatine kinase and **(C)** forelimb grip strength. H&E staining assessed the **(D)** unhealthy/healthy tissue ratio (black arrows) and **(E)** centronucleated fibre proportion (orange arrows) of the quadriceps **(F-H)** and active cardiomyocyte degeneration in **(I)** hearts. Pan macrophage marker, CD68, and Masson’s trichrome staining assessed **(J-M)** immune infiltration and **(N-Q)** fibrosis (orange arrows) of quadriceps and **(R)** heart. **(S)** Mitochondrial oxygen consumption was measured in FDB fibres and **(T)** metabolic phenotypes in response to chemical uncoupling are shown. Extent of DMD phenotype was calculated (*mdx* AGG/*mdx* SED*100) to indicate extent of disease aggravation (17). Data in **B-E, I-J**, **N** and **R** are presented as mean ± SEM and *n* are indicated by individual data points. Panel **S** *n*: WT SED = 7, *mdx* SED = 12, *mdx* AGG = 6. Panel **T** *n*: WT SED = 9, *mdx* SED = 11, *mdx* AGG = 8. Statistical analysis used one-way ANOVA: \**P* <0.05, \*\**P* <0.01, \*\*\**P* <0.001, \*\*\*\**P* <0.0001. **F-H**, **K-M** and **O-Q** scale bar = 50μm.

Exercise aggravation significantly increased plasma creatine kinase (CK) levels in *mdx* mice (Figure 1B) but had no impact on forelimb grip strength (Figure 1C) or other functional measures (Supplemental Figure 1A-C). While neither anthropometric measures nor respiratory or *ex vivo* force deficits were observed (Supplemental Figure 1D-K), aggravation did exacerbate dystrophic histopathology. We analysed quadriceps, which are particularly recruited and damaged during exercise (18, 20), and heart, to assess indices of muscle damage, inflammation, regeneration, immune cell infiltration and fibrosis. Aggravation increased the unhealthy/healthy tissue ratio (*p*=0.0527; Figure 1D) and percentage of regenerating centronucleated fibres indicating greater muscle damage and repair (Figure 1E-H), as well as CD68+ macrophage infiltration (*p*=0.0527; Figure 1J-M) and mature fibrosis (Figure 1N-Q) in *mdx* quadriceps. Aggravation, however, did not affect active cardiomyocyte degeneration and fibrosis of the heart (Figure 1I and R) relative to *mdx* sedentary controls. Perilipin+ adiposis of quadriceps muscle was not impacted by aggravation nor significantly different from WT (Supplemental Figure 1L). We also assessed mitochondrial respiration in flexor digitorum brevis (FDB) fibres using extracellular flux and a mitochondrial stress test (Figure 1S). *Mdx* aggravated FDB fibres had an overall higher energetical state (Figure 1T) in line with increased mechanical damage and regenerative activity relative to sedentary *mdx* and WT controls. However, aggravation had no effect on urinary oxidative stress biomarker, 8-isoprostane (Supplemental Figure 1M).

### DMF improves grip strength and transcriptional control of myofibril assembly

DMF treatment increased forelimb grip strength (corrected for body weight) at the experimental endpoint (Figure 2A) (notably, this measure was higher in *mdx* vehicle mice compared to WT (Figure 1C)) but there was no effect of treatment on the hang test minimal holding impulse, which assesses both strength and endurance (Figure 2B). DMF had no effect on rotarod performance whilst PRED significantly improved time to fall, a measure of neuromotor coordination (Figure 2C). DMF treated alone and in the DMF+PRED combination reduced ambulatory distance whilst PRED had no effect (assessed over 10-minutes in the open-field; Figure 2D). Our functional data suggest that DMF might be better for preserving strength-based measures whereas PRED may be better for endurance.

**Figure 2.**
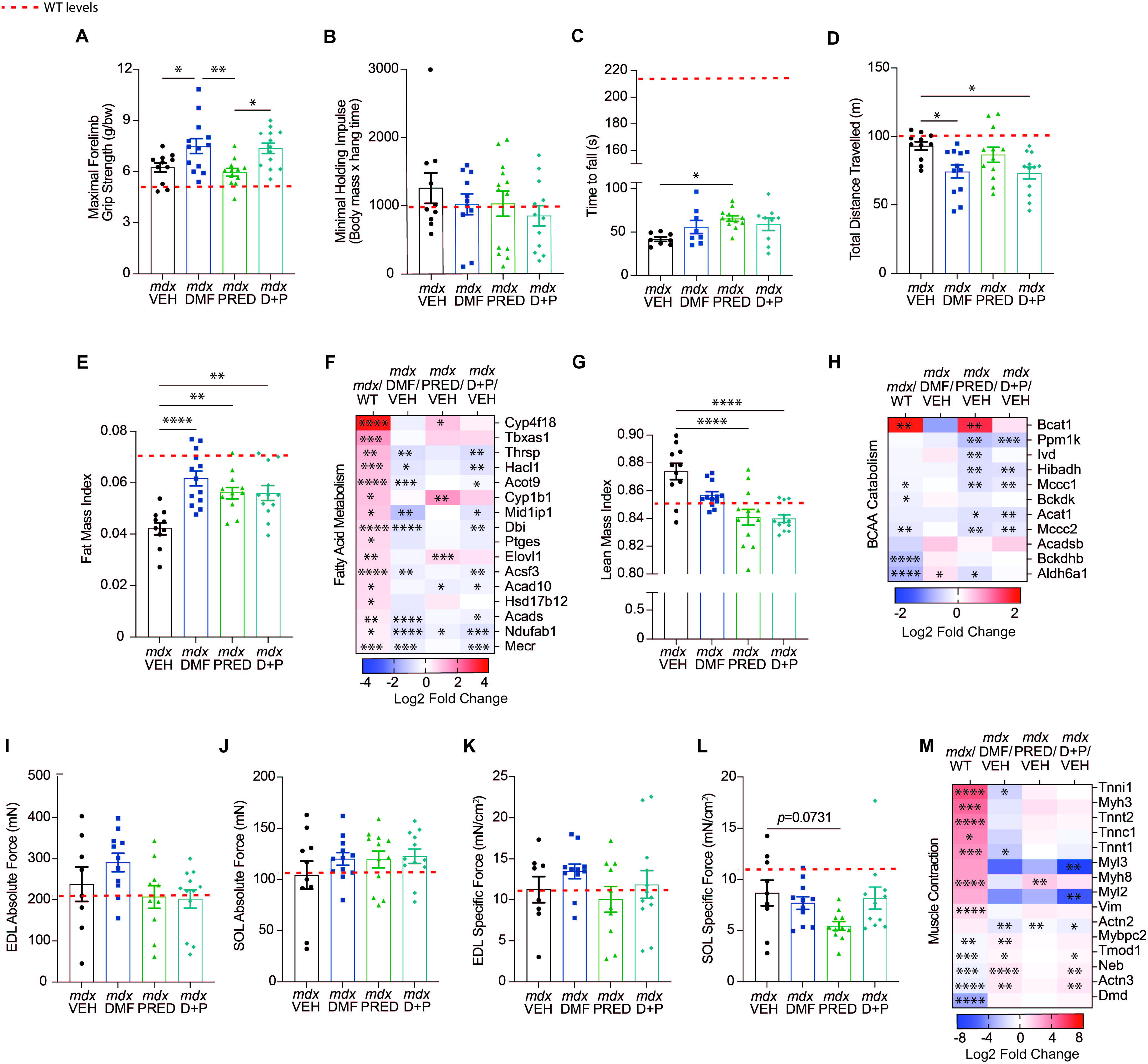
DMF treatment improves grip strength and modulates transcriptional control of myofibril assembly. Functional parameters including **(A)** maximal forelimb grip strength, **(B)** minimal holding impulse (whole body grip strength), **(C)** rotarod time to fall and **(D)** distance travelled in the open field were assessed at the experimental endpoint. Anthropometric measures and their associated transcriptional pathways in muscle were also assessed: **(E)** Fat mass index and **(F)** the fatty acid metabolism pathway, **(G)** lean mass index, and **(H)** the BCAA catabolism pathway. Absolute force was measured *ex vivo* in **(I)** EDL and **(J)** SOL and specific force was calculated **(K-L)**. The transcriptomics dataset was probed for **(M)** muscle contractile associated genes. Data in **F, H** and **M** are based on log2 fold change from WT for *mdx* VEH and *mdx* VEH for treatment groups (DMF, PRED and DMF+PRED). Data in **A-E**, **G**, and **I-L** are presented as mean ± SEM and *n* are indicated by individual data points. Statistical significance was tested via one-way ANOVA: \**P* <0.05, \*\**P* <0.01, \*\*\**P* <0.001, \*\*\*\**P* <0.0001.

In comparison to *mdx* vehicle, all treatments increased the fat mass index (Figure 2E), which was reduced by ~40% in *mdx* relative to WT mice (Supplemental Figure 1D). Accordingly, we probed the muscle (quadriceps) transcriptome for genes associated with fatty acid metabolism. Of the 16 upregulated genes in *mdx* muscle (compared to WT), DMF downregulated 9 genes, PRED downregulated 2 genes and DMF+PRED downregulated 10 genes (Figure 2F). Consistent with the known suppressive effects of PRED on muscle and bone growth (11, 21, 22), PRED treated alone and in the DMF+PRED combination, significantly reduced the lean mass index relative to *mdx* vehicle (Figure 2G). At the muscle transcriptome level, PRED significantly downregulated many genes involved in branched-chain amino acid (BCAA) catabolism but upregulated *Bcat1* (branched chain amino acid transaminase 1), which was already upregulated by 4-fold in *mdx* aggravated relative to WT mice (Figure 2H). PRED reduced liver mass and all treatments normalised the higher quadriceps mass noted in *mdx* aggravated (and sedentary) mice (Supplemental Table 1 and 2) suggesting multi-tissue drivers of lean mass reduction.

At the experimental endpoint (8 weeks of age), extensor digitorum longus (EDL) and soleus muscles, predominantly fast- and slow-twitch respectively, were harvested for *ex vivo* contractile testing. There was no effect of treatments on either EDL or soleus absolute or specific force production (Figure 2I-L) and only DMF+PRED combination affected the force frequency relationship by increasing force production at 60 and 80Hz in the EDL (Supplemental Figure 2A-B). No treatment impacted fatigue or recovery measures (Supplemental Figure 2C-D). Notably, *ex vivo* contraction measures were comparable between WT and *mdx* EDL and soleus (Supplemental Figure 1J-K). We further probed transcriptional control of muscle contraction pathways for molecular changes in mixed fibre type muscle (i.e., quadriceps) that may be induced by treatments but not captured through *ex vivo* studies using fibre type exclusive/predominant muscles (i.e., EDL and soleus). Slow troponin isoforms, *Tnni1* and *Tnnt1*, were significantly upregulated in *mdx* aggravated compared to WT muscle (Figure 2M), consistent with previous studies in dystrophic muscle (23). The downregulation of *Actn2* by all treatments suggests concomitant sarcomere remodelling but only DMF and DMF+PRED treatment upregulated expression of *Tmod1, Neb,* and *Actn3* (and *Mybpc2* for DMF treatment), promoters of sarcomere stability (24, 25) in the absence of dystrophin. PRED upregulated *Myh8,* a gene that has been associated with regenerating DMD muscle (23). There was no effect of treatment on plethysmography (respiratory) readouts (Supplemental Figure 2E-G) or kyphosis index (Supplemental Figure 2H-L), which were unaffected by *mdx* aggravation.

### DMF moderates the immune response in *mdx* quadriceps muscle

Persistent activation of the innate immune system leads to hyperinflammation, oxidative stress and tissue damage in dystrophic muscles (10). The unhealthy/healthy tissue ratio was significantly higher in *mdx* aggravated quadriceps (Figure 1D) but was not reduced by any drug treatment (Figure 3A, C-F). However, there was a trend (*p*=0.0734) for DMF to reduce the percentage of centronucleated regenerating fibres (Figure 3B-F) suggesting modulation of degeneration/regeneration processes. Further investigation of the myogenic transcriptional pathway revealed that *Myog, Myod1, Tcf4, Mapk11* were upregulated in mdx aggravated muscle. DMF downregulated *Myog* and *Mapk11*, PRED downregulated *Myod1 and Mapk12,* and DMF+PRED combination downregulated *Tcf4* highlighting unique modulatory effects. Pan macrophage marker, CD68, was significantly reduced by DMF and PRED (Figure 3H-L) consistent with their anti-inflammatory profiles. Interestingly, combined delivery of DMF+PRED had no effect on macrophage infiltrate. We probed our transcriptomic dataset for common differentially expressed inflammatory markers linked to human DMD, including *Ccl2, Ccl6, Ccl7, Ccl8, Ccr2, Cx3cr1, Nfkb1, Nfkb2,* and *Nfkbib* (Figure 3M). 7 genes were significantly upregulated in *mdx* vehicle quadriceps and DMF treatment significantly downregulated 5 (*Ccl2, Ccl7, Ccl8, Ccr2,* and *Ccl6*) and trended to downregulate *Nfkb2* (*p*=0.0702), a component of the master regulator of innate immunity. Both PRED and DMF+PRED downregulated fewer inflammatory genes: one (*Cx3cr1*) and three (*Ccl7, Ccl2,* and *Ccr2*), respectively.

**Figure 3.**
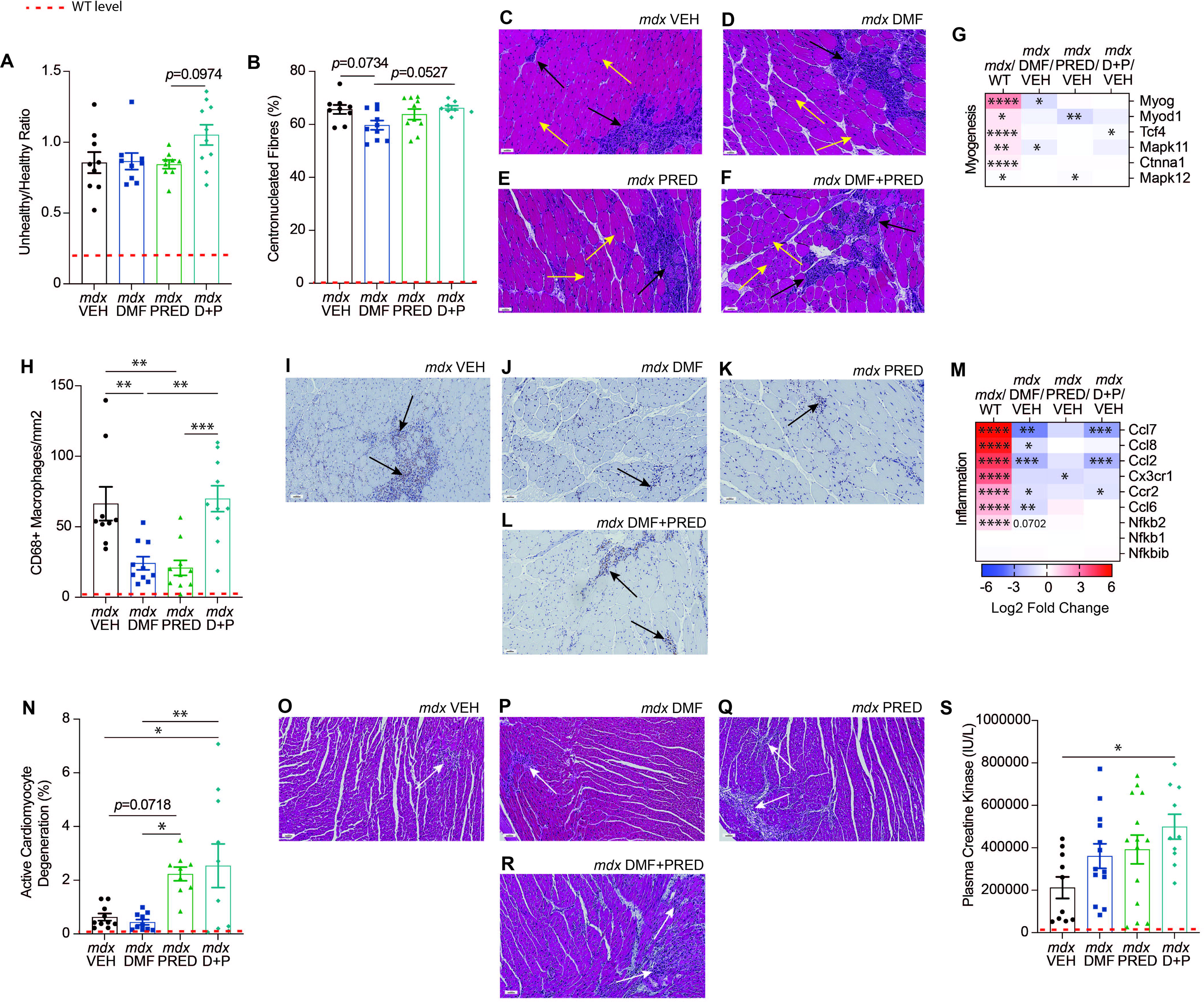
DMF reduces macrophage infiltration of quadriceps. The **(A)** unhealthy to healthy tissue ratio and **(B)** percentage of centronucleated fibres are shown with **(C-F;** yellow arrows indicate centronucleated fibres; black arrows indicate infiltrate) in the quadriceps. **(G)** Transcriptomic signature of myogenesis. **(H)** Pan macrophage marker CD68 was immunohistochemically assessed in the quadriceps and **(I-L)** representative images show CD68+ macrophages (indicated by black arrows). **(M)** The quadriceps transcriptomic dataset was probed for inflammatory markers commonly upregulated in DMD. **(N)** Degenerating area of the heart was also assessed via H&E and **(O-R)** representative images show active cardiomyocyte degeneration (indicated by white arrows). **(S)** Clinically compatible biomarker creatine kinase was assessed at the experimental endpoint. Data in heatmaps (**G** and **M**) are based on log2 fold change from WT for *mdx* VEH and *mdx* VEH for treatment groups (DMF, PRED and DMF+PRED). Data in **A-B**, **H**, **N** and **S** are presented as mean ± SEM and *n* are indicated by individual data points. Statistical significance was tested via one-way ANOVA: \**P* <0.05, \*\**P* <0.01, \*\*\**P* <0.001. **C-F**, **I-L** and **O-R** scale bar = 50μm.

While cardiorespiratory decline is mostly evident in older *mdx* mice, we assessed heart histology since we aggravated our model. Cardiomyocyte degeneration was greater in *mdx* hearts, however, DMF was unable to reduce it. PRED trended to increase cardiomyocyte degeneration (*p*=0.0718) and when combined DMF+PRED significantly increased this histopathologic feature (Figure 3N-R). Endpoint plasma CK levels were unchanged by DMF or PRED treatment but combinatorial DMF+PRED increased CK levels (Figure 3S) suggesting cardiac origin.

### DMF decreases fibrosis in the quadriceps and heart and downregulates key disease driving gene, *Timp1*

Fibrosis is a complex pathology that can result from increased ECM synthesis and/or increased ECM degradation. In a recent meta-analysis, five genes were identified as candidate seed genes driving the DMD disease module (*Timp1, Spp1, Fn1, Mmp2,* and *Igf1*) (26). A more recent study identified a *Bmp4-*induced molecular signature in DMD patient muscles involving upregulated expression of *Serping1, Adamts3, HCAR2, Smad8* and *Unc13c* (27). We show that *Timp1, Spp1,* and *Fn1* (Figure 4A) and *Serping1* (Figure 4B) are upregulated in *mdx* aggravated quadriceps alongside fibrosis (Figure 4C-D). DMF treatment significantly decreased *Timp1* (inhibitor of matrix metalloproteinases (MMPs)) expression (Figure 4A) and fibrosis of quadriceps (Figure 4C and E). In contrast, PRED upregulated *Spp1, Fn1*, *Igf1* (all notably upregulated by glucocorticoids (28–30)) and *Bmp4* transcription and in combination DMF+PRED increased *Spp1* expression only (Figure 4A). PRED alone had no effect on fibrosis in the quadriceps however, DMF+PRED treatment worsened it (Figure 4C, F-G). We probed 3 Reactome-based fibrosis pathways in our transcriptomics dataset: collagen formation, ECM degradation and ECM organisation. Overall, DMF and PRED induced very different ECM signatures (Figure 4H-J). PRED appeared to upregulate collagen formation (Figure 4H), degradation and organisational gene expression suggesting functional remodelling although not pathological fibrosis per se. DMF was the only treatment to reduce cardiac fibrosis (Figure 4K-O) consistent with its anti-fibrotic effects in quadriceps (Figure 4C).

**Figure 4.**
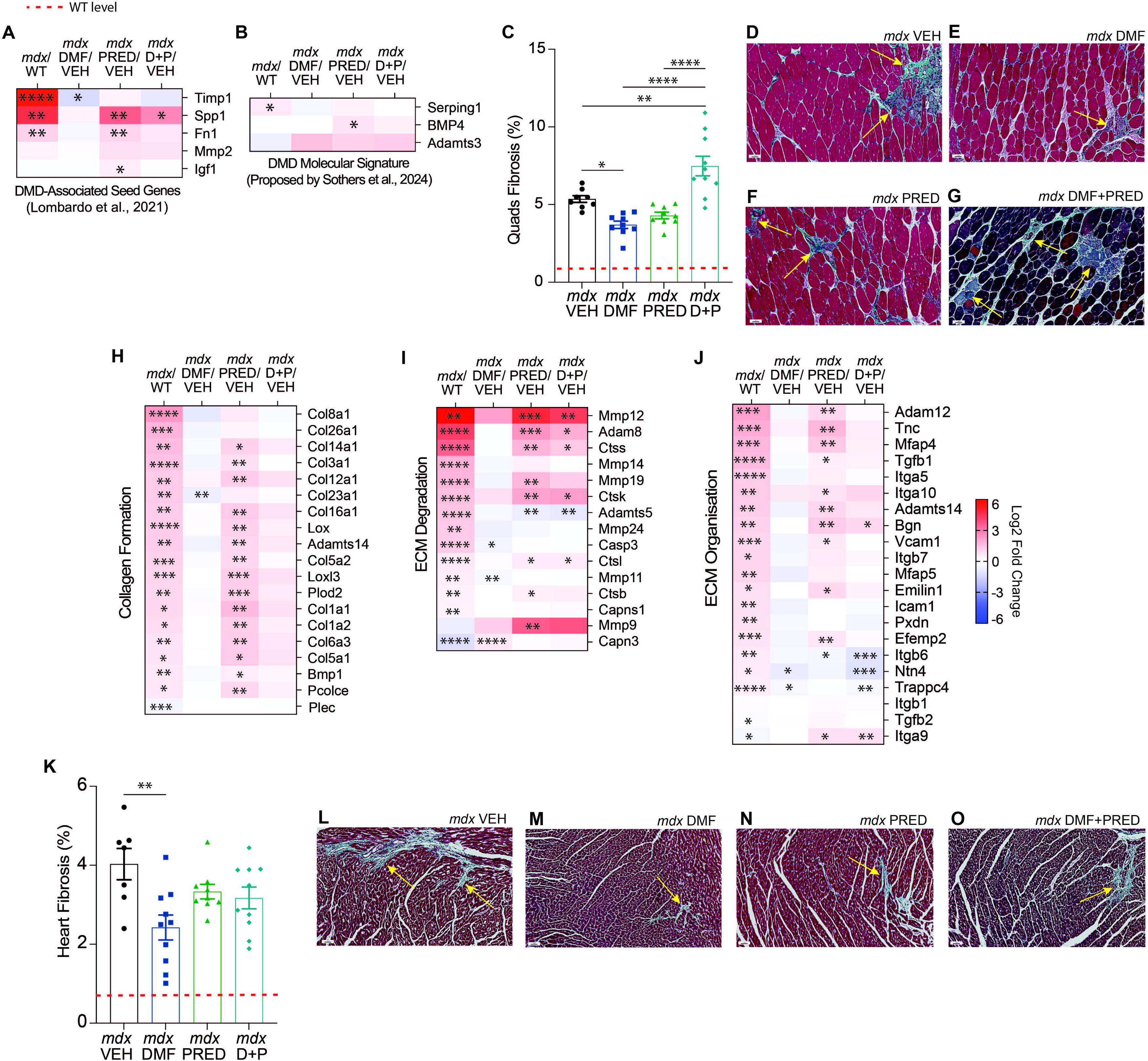
DMF reduces fibrosis of quadriceps and heart. Expression of the proposed **(A)** fibrosis-associated DMD seed genes (26) and **(B)** DMD molecular signature (27) was probed using our transcriptomic dataset. **(C)** Quadriceps fibrosis was quantified using Masson’s trichrome staining and **(D-G)** representative images are shown. Transcriptional pathways related to **(H)** collagen formation, **(I)** ECM degradation and **(J)** ECM organisation were analysed to investigate mechanisms of anti-fibrosis. **(K)** Fibrotic area of the heart was also assessed and **(L-O)** representative images are shown. Data in all heatmaps **(A-B** and **H-J)** are based on log2 fold change from WT for *mdx* VEH and *mdx* VEH for treatment groups (DMF, PRED and DMF+PRED). Data in **C** and **K** are presented as mean ± SEM and *n* are indicated by individual data points. Statistical significance was tested via one-way ANOVA: \**P* <0.05, \*\**P* <0.01, \*\*\**P* <0.001, \*\*\*\**P* <0.0001. **D-G** and **L-O** scale bar = 50μm.

### DMF reduces muscle adiposis through modulation of the adipogenic and sphingolipid metabolism transcription programs in *mdx* muscle

Replacement of muscle fibres with adipose tissue is prominent in human DMD muscle but minimal in *mdx* muscle, i.e., perilipin-1, a protein that localises to adipocytes, was comparable between WT and *mdx* quadriceps (Supplemental Figure 1L). Nevertheless, DMF treatment decreased adipocyte abundance whereas PRED, alone and in combination with DMF, had no effect (Figure 5A-E). To interrogate the mechanisms, we probed lipid metabolism and fibro-adipogenic progenitor (FAP)-related pathways in our transcriptomics dataset. Of the 3 established FAP-associated marker genes in humans, (*Cd34, Pdgfra,* and *Dcn*), *Cd34* expression was upregulated in *mdx* quadriceps and was downregulated by DMF (Figure 5F). Reactome pathway analysis revealed that the tumour necrosis factor receptor (TNFR)-mediated ceramide production pathway was significantly upregulated in *mdx* quadriceps and was normalised by DMF but not PRED treatment (Figure 8I). We probed genes within this pathway (Figure 5G) and revealed DMF normalised *Tnfrsf1a* (a pro-inflammatory biomarker in human DMD (31)) and *Nsmaf* expression. Sphingolipid metabolism pathways were also probed since DMF is known to modulate sphingolipids through HCAR2 agonism in the context of MS (32). Of the 11 upregulated sphingolipid metabolism genes identified in *mdx* muscle, DMF downregulated 2 while PRED and DMF+PRED significantly upregulated 9 and 5 genes, respectively (Figure 5H). Furthermore, DMF normalised the expression of *Sgms1* and *Cers1*, which were downregulated in *mdx* quadriceps. 10 adipogenesis genes were differentially expressed in *mdx* muscle: 4 upregulated and 6 downregulated (Figure 5I). DMF was more effective at moderating the adipogenic gene program than PRED, either alone or combined. Finally, we used targeted lipidomics to assess the lipid species profile in quadriceps muscles. 20 differentially expressed lipid species were detected in *mdx* quadriceps relative to WT (11 upregulated and 9 downregulated), but these were not modulated by any treatment (Figure 5J). Of note, both PRED and DMF+PRED modulated 1 and 4 normally expressed lipid species, respectively.

**Figure 5.**
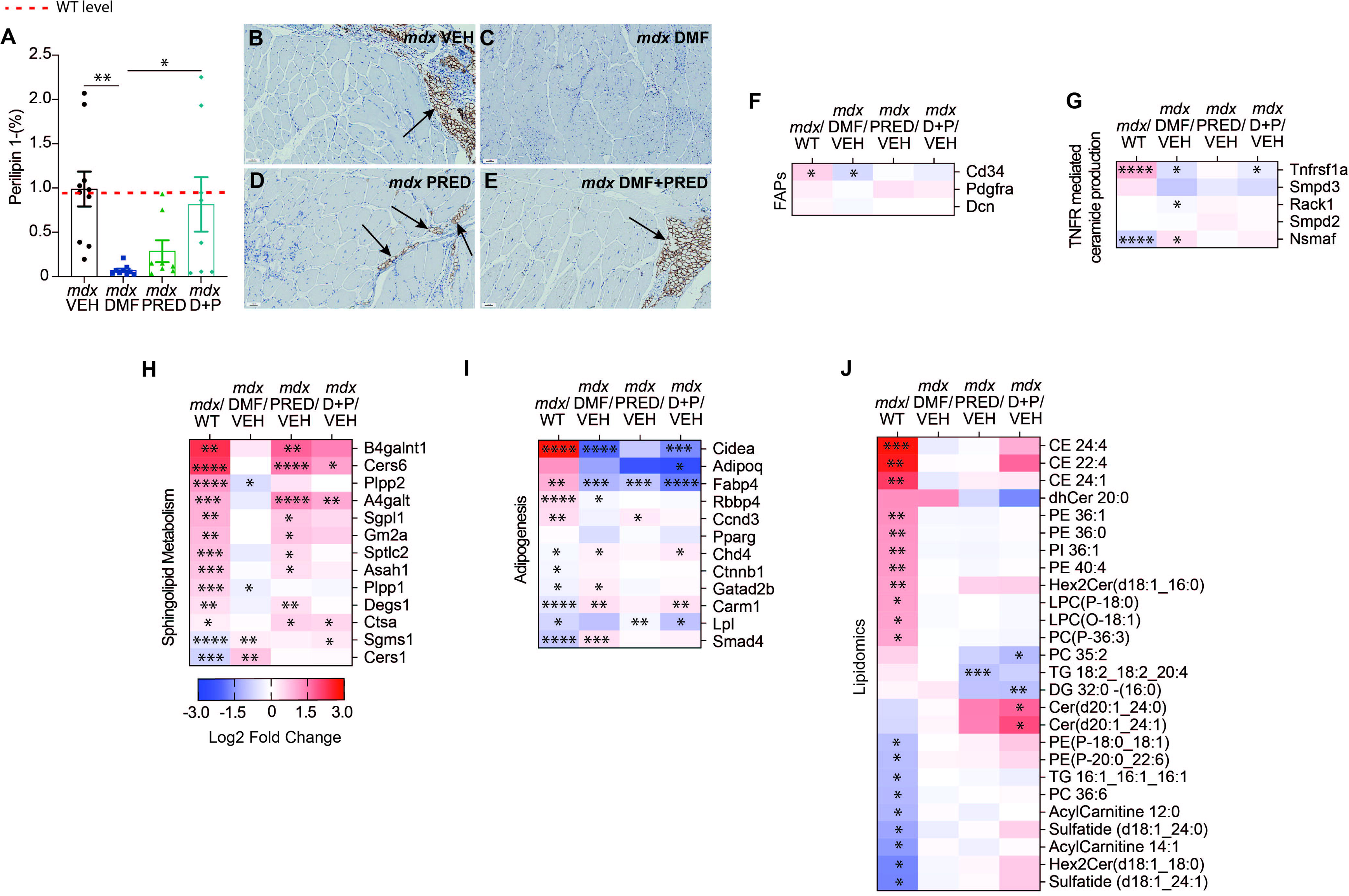
DMF decreases muscle lipid content through modulation of adipogenesis and sphingolipid metabolism pathways in quadriceps. **(A-E)** Percentage of perilipin-1+ lipid droplets in quadriceps. Lipid associated transcriptional pathways were probed including **(F)** FAPs, **(G)** TNFR mediated ceramide production, which was a significantly altered Reactome pathway in *mdx* aggravated relative to WT quadriceps, **(H)** sphingolipid metabolism and **(I)** adipogenesis. **(J)** Targeted lipidomics was performed to reveal the relative abundance of muscle lipid species. Data in all heatmaps **(F-J)** are based on log2 fold change from WT for *mdx* VEH and *mdx* VEH for treatment groups (DMF, PRED and DMF+PRED). Data in **A** is presented as mean ± SEM and *n* are indicated by individual data points. Statistical significance was tested via one-way ANOVA: \**P* <0.05, \*\**P* <0.01, \*\*\**P* <0.001, \*\*\*\**P* <0.0001. **B-E** Scale bar = 50μm.

### DMF significantly recovers inflammatory, fibrosis and fat mass associated parameters

The recovery score is used to express the effect of a treatment, not just by the difference between treated and untreated *mdx* mice, but relative to the extent of the deficiency between WT and *mdx* mice (17). We calculated the recovery score (Figure 6) of parameters that were significantly affected by murine DMD aggravation. CD68 macrophage infiltration of quadriceps was improved by both DMF and PRED to a similar extent (65% vs. 66%, respectively) whilst DMF+PRED showed no improvement (0.78%). Fat mass index was recovered more so by DMF and DMF+PRED (59% and 44%, respectively) than PRED, which had no effect (−1.27%). DMF was the most effective treatment at recovering cardiac fibrosis (46%) followed by PRED and DMF+PRED, which had a comparable recovery score (20% and 24%, respectively). Similarly, DMF was the most effective treatment to recover quadriceps fibrosis (37%) followed by PRED (23%). DMF+PRED worsened quadriceps fibrosis resulting in a negative recovery score (−47%).

**Figure 6.**
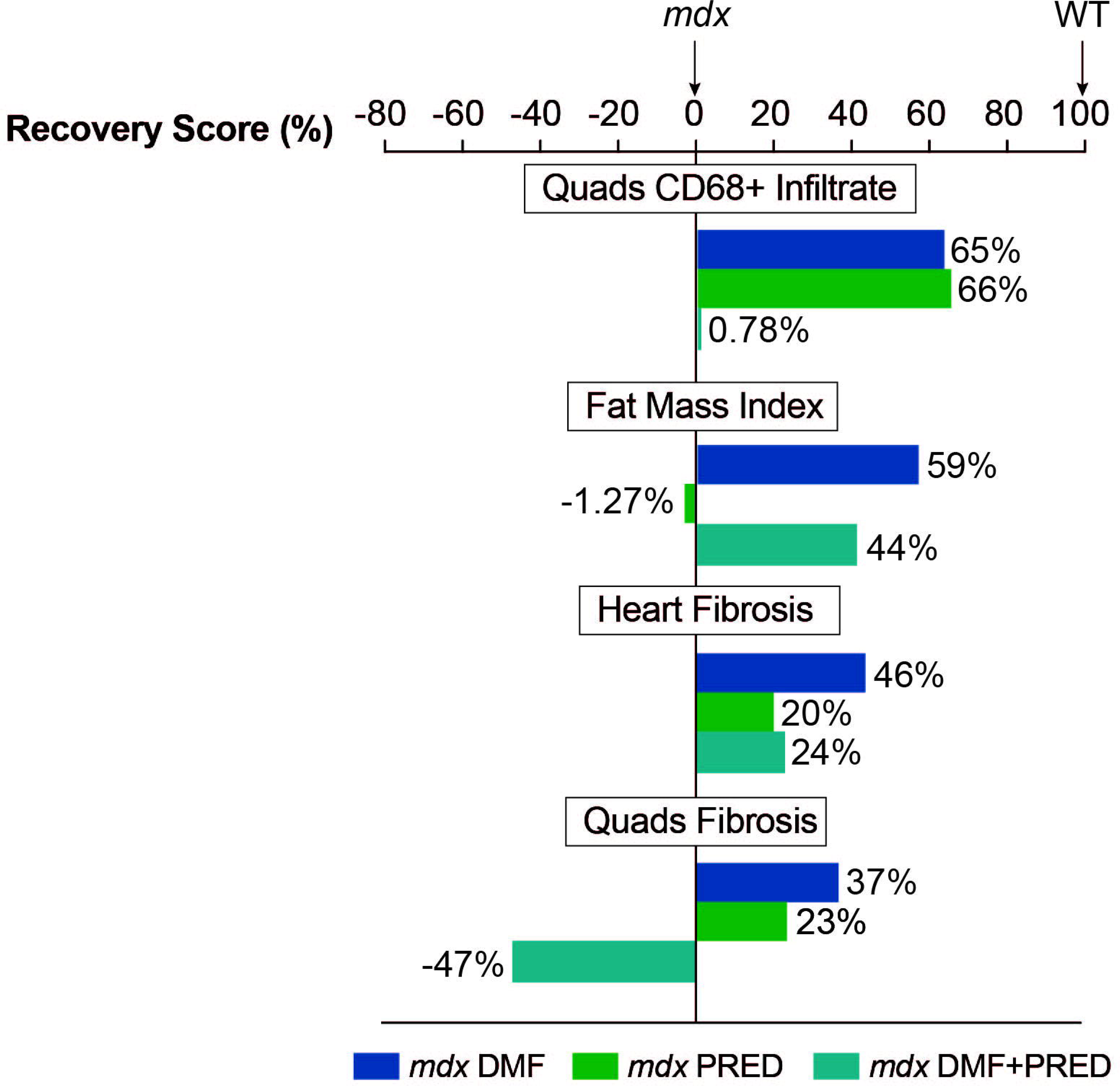
The impact of the treatments on the *mdx* mouse. The recovery score is used to express the effect of a treatment, not just by the difference between treated and untreated *mdx* mice, but relative to the extent of the deficiency between WT and *mdx* mice. A recovery score of 100% indicates the parameter is equal to that of the WT while a score of 0% indicates no improvement has been made. This was calculated in accordance with TREAT-NMD SOP (DMD_M.1.1_001) (89) as follows: (*mdx* treated)-(*mdx* untreated)/(WT)-(*mdx* untreated) x 100.

### DMF transcribes a highly specific anti-inflammatory and anti-oxidation signature in muscle

A well-established MOA of DMF is via activation of Nrf2, which initiates a potent cytoprotective response (Figure 7A). Deep transcriptome profiling did not reveal changes to expression of *Nf2el2* (Nrf2), or its negative repressor, *Keap1*, in response to DMF treatment. However, DMF normalised the otherwise upregulated expression of proteasomal subunit genes, *Psmb1, Psme1, Psma4,* and *Psma1* (Figure 7B), which degrade Nrf2 and prevent its activity. DMF also tempered expression of antioxidant/detoxification enzymes *Ccs, Prdx2,* and *Txn2* (Figure 7C). Our pathways analysis revealed DMF trended to drive DNA damage reversal pathways (FDR=0.0686; data not shown), thus we further probed this pathway. While *Alkbh5* was the only downregulated gene in *mdx* aggravated quadriceps, DMF upregulated the expression of both *Alkbh5* and *Fto*, and DMF+PRED upregulated *Alkbh5* expression (Figure 7D). We next probed the interleukin (IL) signalling pathways where 15 genes were differentially expressed in *mdx* quadriceps (14 upregulated, 1 downregulated; Figure 7E). Of the 14 upregulated genes, DMF reduced the expression of 5, PRED further upregulated 3 whereas DMF+PRED up- and down-regulated 2 each. DMF also modulated more differentially expressed genes associated with the TNF signalling pathway than either PRED or DMF+PRED (Figure 7F). DMF reduced 4 upregulated genes related to the pro-inflammatory response (*Fadd, Tnfrsf1a, Otulin,* and *Rps27a*) and upregulated 5 genes associated with anti-inflammation (*Cyld, Usp4, Spata2, Sppl2a,* and *Xiap*). Comparatively, PRED further downregulated *Optn* expression and DMF+PRED normalised *Tnfrsf1a*. Treatments had no effect on urinary oxidative stress biomarker, 8-isoprostane (Figure 7G), albeit 8-isoprostane levels were comparable between *mdx* and WT samples (Supplemental Figure 1M).

**Figure 7.**
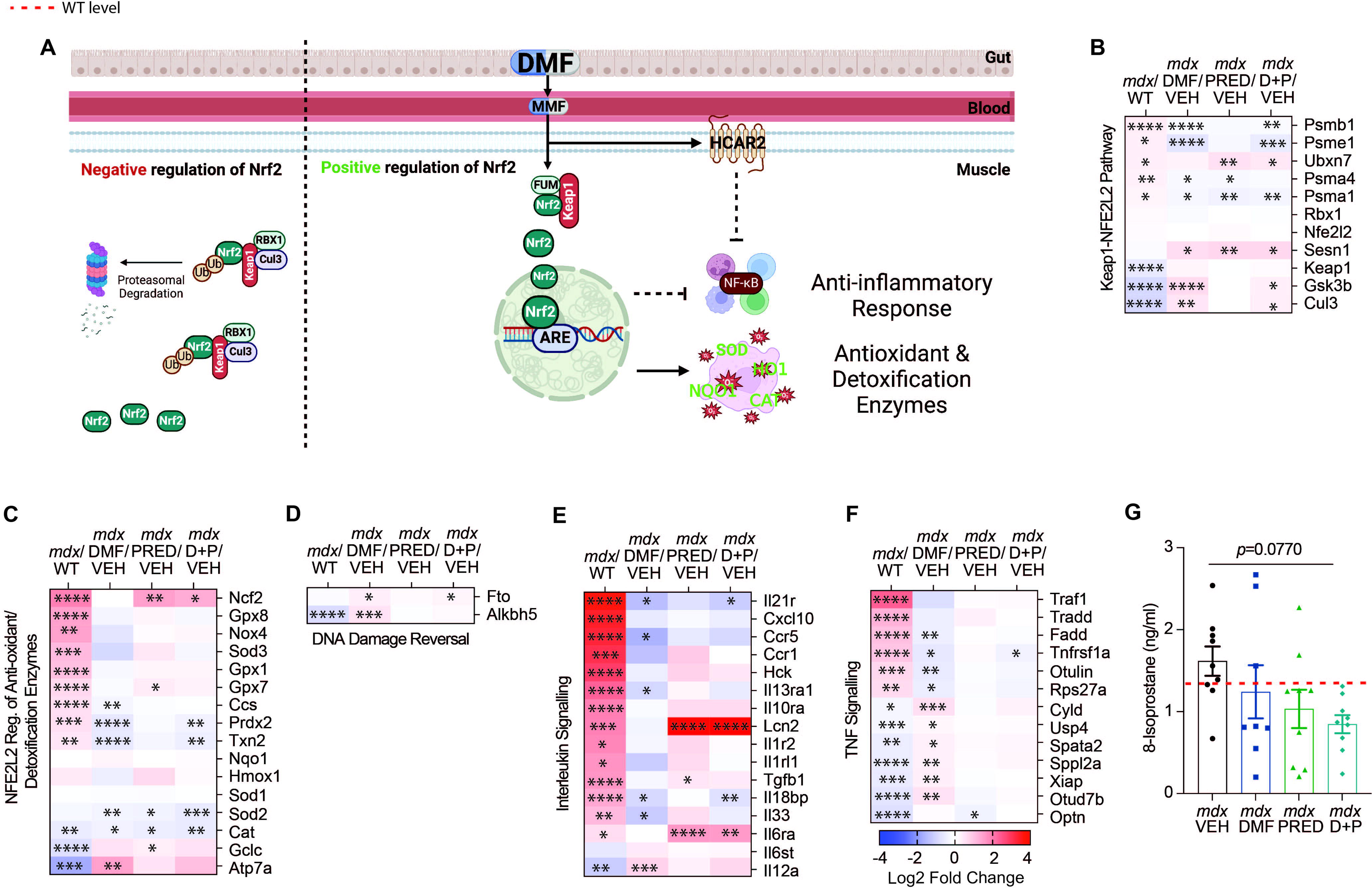
DMF treatment tempers expression of detoxification enzymes and transcribes a highly specific anti-inflammatory signature in muscle. **(A)** DMFs MOA involves dual anti-inflammatory and -oxidative function through transcription of Nrf2. The muscle transcriptome was probed for molecular pathways involved in DMF’s MOA: **(B)** Keap1-Nrf2, **(C)** Nrf2 regulation of antioxidant/detoxification enzymes and **(D)** DNA damage reversal pathways. DMF’s anti-inflammatory capacity was investigated through **(E)** interleukin and **(F)** TNF signalling pathway analysis, and urinary oxidative stress marker **(G)** 8-isoprostane. All heatmap data **(B-G)** are based on log2 fold change from WT for *mdx* VEH and *mdx* VEH for treatment groups (DMF, PRED and DMF+PRED). Data in **H** are presented as mean ± SEM and *n* are indicated by individual data points. Statistical significance was tested via one-way ANOVA: \**P* <0.05, \*\**P* <0.01, \*\*\**P* <0.001, \*\*\*\**P* <0.0001.

### DMF reduces purine degradation and mitobiogenic stimuli but has no effect on respiratory flexibility

Since short-term DMF increased mitochondrial respiration via anaplerosis (16), we assessed metabolic function in animals via a fatigue run and respiratory parameters in response to moderate-term treatment (Figure 8A). DMF had no significant effect on fatigability during running (Supplemental Figure 3A) or basal, ATP-linked, maximal, or non-mitochondrial respiration (Supplemental Figure 3B-E). PRED decreased coupling efficiency (Supplemental Figure 3F) and spare respiratory capacity (Supplemental Figure 3G), a measure of mitochondrial responsivity to stress, while DMF+PRED reduced maximal respiration and the spare respiratory capacity. Metabolic phenograms were produced to survey proportionate oxidative (OCR) to glycolytic (ECAR) metabolism and revealed *mdx* vehicle FDB fibres were more energetic – an effect that was normalised by all treatments (Figure 8B).

**Figure 8.**
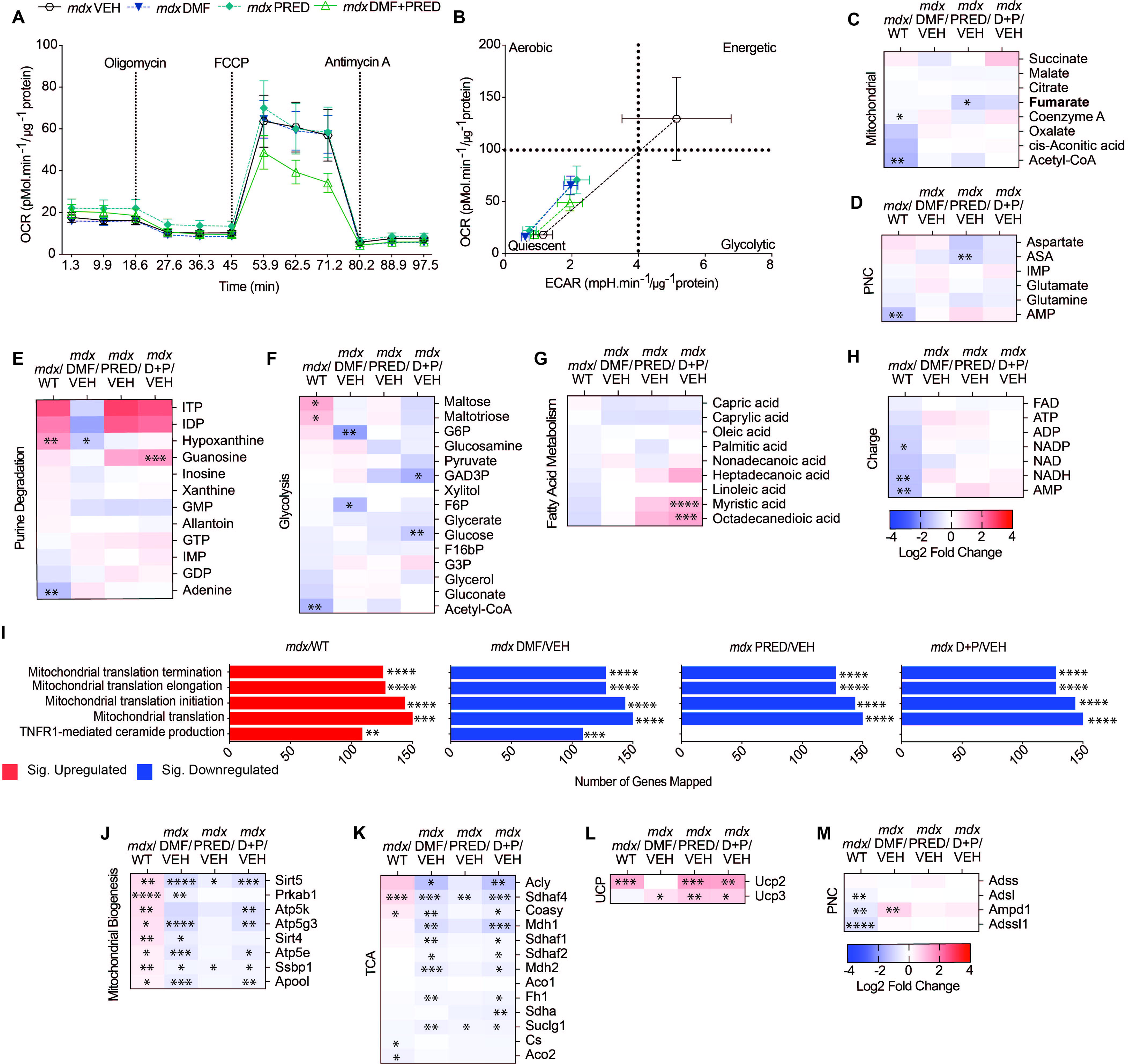
DMF reduces purine degradation and mitobiogenic stimuli. **(A)** Mitochondrial oxygen consumption was measured in FDB fibres and **(B)** metabolic phenotypes in response to chemical uncoupling are shown. Untargeted polar metabolomics was performed on quadriceps muscle and **(C)** mitochondrial, **(D)** PNC, **(E)** purine degradation, **(F)** glycolysis, **(G)** fatty acid metabolism, and **(H)** charge metabolomes are presented. **(I)** Top 5 altered Reactome pathways of the transcriptomics dataset. Reactome pathways associated with **(J)** mitochondrial biogenesis, **(K)** TCA cycle, **(L)** UCP and **(M)** PNC. Data in all heatmaps **C-H** and **J-M** are based on log2 fold change from WT for *mdx* VEH and *mdx* VEH for treatment groups (DMF, PRED and DMF+PRED). Panel **A** *n*: *mdx* VEH = 6, *mdx* DMF = 12, *mdx* PRED = 12, *mdx* DMF+PRED = 12. Panel **B** *n*: *mdx* VEH = 8, *mdx* DMF = 12, *mdx* PRED = 12, *mdx* DMF+PRED = 12. Statistical significance was tested via one-way ANOVA: \**P* <0.05, \*\**P* <0.01, \*\*\**P* <0.001, \*\*\*\**P* <0.0001.

Untargeted polar metabolomics revealed reduced mitochondrial tricarboxylic acid (TCA) cycle (Figure 8C), purine nucleotide cycle (Figure 8D), purine nucleobase (Figure 8E), charge/energy transfer (Figure 8H) and pyrimidine (Supplemental Figure 3I) metabolites in *mdx* aggravated quadriceps relative to vehicle controls, indicating a stimulus for hyper-energetic metabolism. Purine degradation and excretion metabolite, hypoxanthine, was increased (Figure 8E), along with glycolysis associated sugars, maltose and maltotriose (Figure 8F), were increased in *mdx* aggravated quadriceps suggesting metabolic stress. No differences in fatty acid (Figure 8G) or polyamine metabolism (Supplemental Figure 3H) were observed. Of the 10 altered metabolites, DMF normalised hypoxanthine. PRED and DMF+PRED combination had no effect. The full metabolomic signature is presented in Supplemental Figure 3J and principal component analysis (PCA) and volcano plots are shown in Supplemental Figure 4.

Transcriptomic pathways analysis revealed pathways associated with mitochondrial termination, elongation, initiation, and translation and TNFR1-mediated ceramide production were upregulated in *mdx* muscle (Figure 8I and Table 1). All treatments downregulated these pathways, except for the TNFR1-mediated ceramide production pathway, which was only downregulated by DMF. We probed our dataset to see whether stress-induced mitochondrial biogenesis transcription could explain our pathways analysis (Figure 8J). Eight mitochondrial biogenesis genes were upregulated, but transcription of traditional biogenesis enzyme biomarker, *Cs* (citrate synthase) was reduced in *mdx* muscle (Figure 8K). DMF and DMF+PRED downregulated more mitochondrial biogenesis genes than PRED (7 vs. 6 vs. 2, respectively). Two uncoupling proteins (UCP) genes were probed: *Ucp2* was significantly upregulated in *mdx* aggravated quadriceps and further upregulated by PRED and DMF+PRED treatment, while *Ucp3* was normally expressed in *mdx* muscle, but upregulated by all 3 treatments (Figure 8L). Since fumarate is also endogenously produced by the purine nucleotide cycle (PNC), we probed this pathway for treatment effects. All 3 PNC genes were downregulated in *mdx* quadriceps. DMF upregulated *Ampd1* (Figure 8M).

**Table 1.**
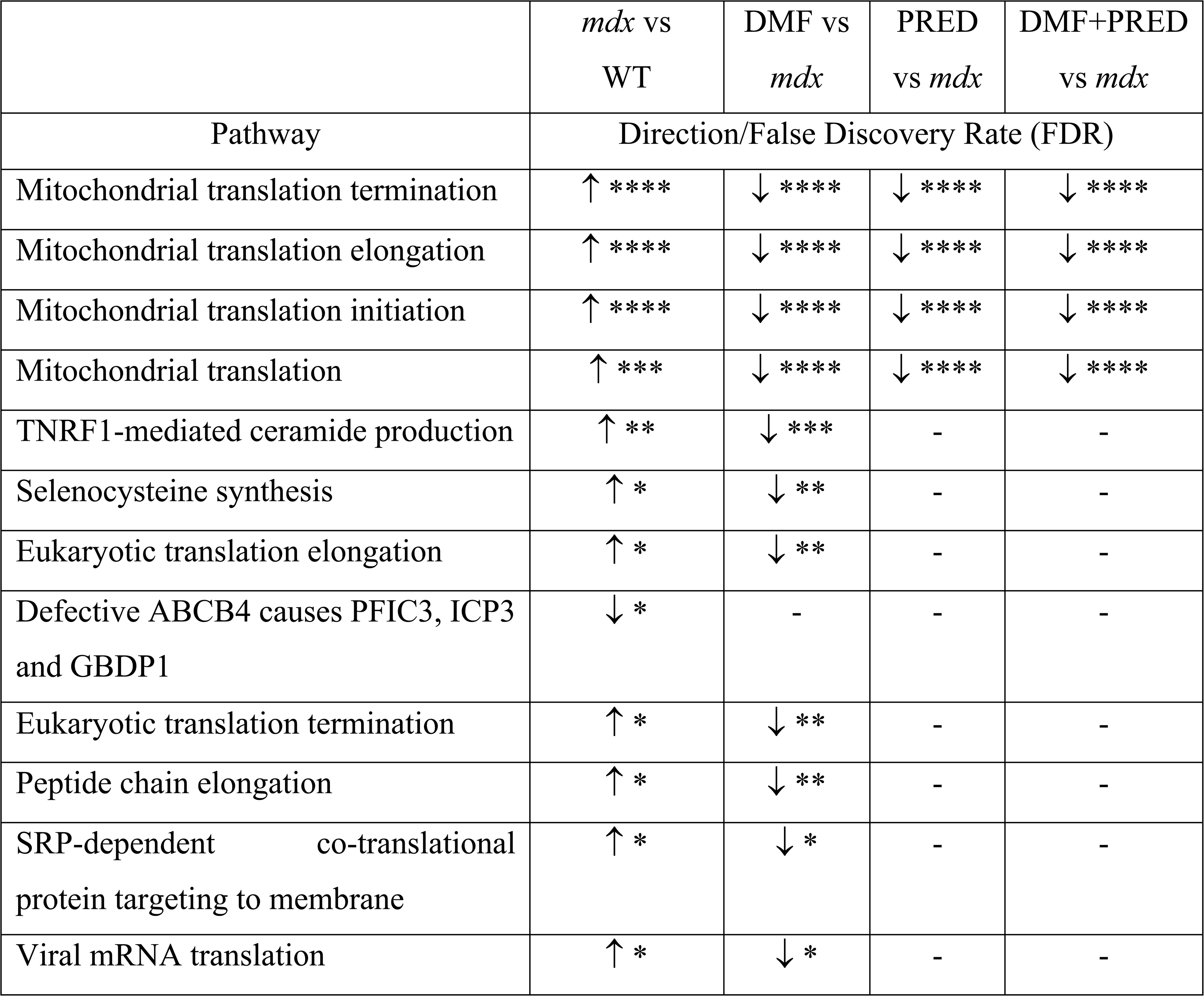
Effect of DMF, PRED and DMF+PRED treatment on significantly differentially expressed pathways in *mdx* vs WT. \**P* <0.05, \*\**P* <0.01, \*\*\**P* <0.001, \*\*\*\**P* <0.0001.

## Discussion

Our methodological approach of twice weekly forced treadmill running was successful to increase clinically relevant muscle damage biomarker, plasma CK, and histopathological hallmarks in hindlimb quadriceps muscle. Cardiorespiratory phenotypes are usually only evident from ~12-18 months age in *mdx* mice (33) and it was encouraging that cardiac muscle degeneration and fibrosis were induced by exercise aggravation. Increasing the frequency (e.g., thrice weekly running), duration (e.g., >20 mins per session) or difficulty (e.g., downhill running) of the exercise regimen could further induce both histopathological and functional phenotypes across all impacted physiological systems in future work.

Our data highlight that DMF’s MOA over the moderate term involves highly specific immune modulation, possibly via HCAR2 agonism. We show that DMF was able to recover histopathological exacerbations induced by aggravation of the *mdx* phenotype by on average, 50% (Figure 6). In contrast, PRED recovered on average, 26%. DMF is twice as effective at treating these measures and from a safety standpoint could be better, since it does not reduce lean tissue mass like PRED. In our short-term DMF screen, we demonstrated Nrf2 activation via increased expression of Phase II antioxidant enzymes, NQO1 and SOD1 (16), the biomarkers of DMF’s MOA and efficacy in MS (34). However, in this study, moderate-term DMF treatment, at least transcriptionally, did not maintain NQO1 or SOD1 upregulation. DMF likely induces temporal Nrf2 activation in response to environmental (oxidant, inflammation) cues to maintain a proportionate antioxidant defence system. For example, GSK-3β (glycogen synthase kinase 3β) phosphorylates Nrf2 resulting in its nuclear exclusion and degradation independent of Keap1 (35). GSK-3β is possibly an oxidant sensitive override mechanism to avoid hyper-Nrf2-activation. DMF treatment (alone and in combination with PRED) increased GSK-3β transcription indicating this mechanism is predominate and antioxidative protein expression sufficient to handle the myocellular redox climate.

The most striking effect of DMF in our study was reducing fibrosis of the heart and quadriceps although it is important to note that fibrosis was mild in our 8-week-old *mdx* mice (~4-5%, consistent with others (36, 37)) even with exercise aggravation. Recently, we showed DMF could modulate fibrosis-associated seed genes (*Mmp2, Spp1,* and *Timp1*) (16) that drive the DMD disease program described by Lombardo *et al.* (26). Here we show that moderate-term DMF treatment sustains *Timp1* suppression, whilst PRED significantly upregulates expression of 3 seed genes (*Spp1, Fn1,* and *Igf1*). DMF downregulated key genes involved in collagen formation, ECM degradation and organisation whilst PRED significantly upregulated over more than half (34 out of 54 differentially expressed genes). DMF was previously shown to ameliorate lung (38–40) and renal (41) fibrosis indicating systemic anti-fibrotic function. Of the several possible molecular mechanisms regulating DMFs anti-fibrotic effect, our data indicate reduced (CD68+) macrophage recruitment/accumulation/transition as central. We saw no evidence of DMF-dependent inhibition of TGF-ß/Smad3 signalling or Spp1/osteopontin expression. Notably, when treated in combination DMF+PRED exacerbated exclusively quadriceps fibrosis (−47% recovery score). This effect could be a result of over-immunosuppression, highlighting the intricate symphony of immunology required to maintain muscle over fibrotic tissue. As a weight-bearing and high force generating muscle group, quadriceps are more prone to damage than the heart and therefore seem more sensitive to the detrimental effect of over-immunosuppression. This aspect requires careful consideration in the planning of future clinical trials testing DMF and other immunomodulatory/anti-fibrotic agents for DMD where SOC may need to be withheld or exclusively steroid naïve participants recruited.

In the context of human MS, DMF is known to re-program fat metabolism, especially sphingolipid metabolism (32). Transcripts of the sphingolipid *de novo* biosynthesis pathway are upregulated in skeletal muscle of patients with DMD as well as in other muscular dystrophies (42). Our data highlight multiple points of control of systemic and muscle-specific adipose tissue content. *Mdx* mice (sedentary and aggravated) are particularly lean (low fat mass index relative to WT) and our multi-omic data point to increased metabolic demand as a mechanism (43). DMF treatment normalised these multi-omic signatures and the fat mass of *mdx* animals. The primary active metabolite of DMF, monomethyl fumarate, is a ligand for HCAR2 which is highly expressed on adipocytes and immune cells (44). HCAR2 agonism prevents lipolysis through inhibition of adipose triglyceride and hormone sensitive lipases (32). In contrast to these systemic effects, DMF reduced adipocyte abundance (perilpin-1) at the muscle level in *mdx*. Via its distinct immunomodulatory effects on macrophage sub-populations, DMF downregulated FAP-associated marker genes, which is important because a subset of FAPs can undergo adipogenic differentiation contributing to fatty infiltration of human DMD muscle (45). DMF also reduced *Tnfrsf1a* transcripts which were upregulated in *mdx* muscle. In humans, TNFR1 agonism contributes to muscle adiposis by inducing sphingomyelin hydrolysis-mediated ceramide production. Ceramides are bioactive molecules capable of activating apoptotic and pro-inflammatory pathways and their accumulation in muscle is linked to multiple myopathies (46, 47). Consistent with a study by Laurila *et al.* (42) in *mdx* mice, we show that the sphingolipid metabolism pathway is upregulated in dystrophic muscle. Although DMF downregulated only 2 of 10 significantly upregulated genes within this pathway, PRED led to further upregulation of 9 genes. Sphingolipid metabolism is heightened in MS resulting in ceramide production (48) and more recently, sphingosine-1-phosphate (S1P), as well as transcripts of other sphingolipids were shown to be highly upregulated in dystrophic human and *mdx* muscle (49). Muscle adiposity is much more of a problem in human than *mdx* DMD (50, 51), as is obesity (52, 53). Whether DMF will be efficient enough in humans given that the adipose pathology is worse is an important translational question for follow up. If it were to be similarly effective against human DMD muscle adiposis, DMF could be a potential candidate for repurposing.

A notable effect of short-term but not moderate-term DMF treatment was mitochondrial anaplerosis (16). TCA metabolites, including fumarate, are reduced in dystrophic muscle (54–56). Endogenous fumarate production is achieved within the mitochondrial TCA cycle via reversible fumarase activity from substrate succinate or malate, and the cytosolic PNC via adenylosuccinate lyase (ADSL) from substrate adenylosuccinate. ADSL-synthesised fumarate can be transported into the mitochondria via the malate-aspartate shuttle (after conversion of fumarate to malate in the cytosol) where exchanged aspartate is redirected into the PNC to synchronise these two metabolic systems (57). Adding exogenous fumarate to the system, i.e. via DMF treatment, at least acutely, amplifies ATP production potential. However, prolonged exposure in the present study, drove transcriptional changes to reduce mitochondrial ATP producing capacity. Historically, purine depletion was linked to DMD pathogenesis, and especially, to deficient purine salvage mechanisms (58). The PNC functions to trap inosine monophosphate (IMP) produced from ATP hydrolysis or *de novo* purine biosynthesis, protecting it from further degradation (IMP>inosine>hypoxanthine>xanthine) and flux from muscle (as uric acid). Our data show transcriptional downregulation of all three PNC enzymes (*Adss1*, *Adsl* and *Ampd1*) in *mdx* muscle as has been reported in human DMD muscle (59), and increased degraded purine metabolites, hypoxanthine, and adenine. DMF treatment normalised *Ampd1* expression and hypoxanthine levels. Mechanistically, defending purine salvage may be a useful strategy to de-burden mitochondria and ultimately to reduce reactive oxygen species production. Mitochondria are important signalling hubs directing cellular metabolism, innate immunity, cell death and the stress response. While mild mitochondrial stress may be beneficial to trigger a protective response (mitohormesis) severe and/or prolonged energy stress may contribute to oxidant-induced damage, fibroblast activation, and ECM pathology (60).

Our study reveals important translational aspects. Firstly, immune system involvement in DMD progression is complex as its pharmacological control. Timely and successful muscle regeneration and repair is highly dependent on a well-regulated inflammatory cascade (61) – chronic inflammation and cytokine signalling can promote macrophage phenotype shifts and muscle pathology, while over-immunosuppression can dysregulate muscle regeneration. Alone, DMF and PRED share similar anti-inflammatory and immunomodulatory capability (transcriptionally) to effectively reduce macrophage infiltration of muscle. From a translational perspective, DMF could be a relevant PRED surrogate to slow DMD progression on this point. However, our omics studies reveal vastly different molecular mechanisms, and it is currently unclear which mechanism provides the best long-term results. While our data indicate DMF is a more effective modulator of fibrosis, it would be remiss not to point out that we used a higher PRED dose (5 mg.kg^-1^/day) than commonly used in pre-clinical studies, yet a clinically precise one (0.75 mg.kg^-1^/day guide-lined human dose (62) corrected for mouse metabolism using 12.3 multiplier applied as per FDA guidelines (63)), which may have over-immunosuppressed mice. When combined, DMF+PRED exacerbated quadriceps fibrosis highlighting the probability of over-immunosuppression, drug interactions or both. Recent protocols for murine PRED dosing have emerged demonstrating pulsed delivery (1 mg.kg^-1^ once per week) best mimics human comparable benefits of daily treatment (64). Similarly, weekend pulsing (1-4 mg.kg^-1^ on both days) maintains clinical benefits but with reduced side-effects in DMD patients (65). While our PRED dosing strategy somewhat limits the translational relevance of our DMF findings based on head-to-head comparison, it does flag the implications of overdoing immunomodulation. Secondly, while DMF did have clear anti-fibrotic and anti-lipogenic properties, we saw no improvement of plasma CK levels and muscle degeneration/regeneration histology, the most impacted *mdx* aggravated variables, or *in vivo* and *ex vivo* muscle function which were remarkably comparable to WT. *Mdx* mice in particular have an increased sensitivity to stress (66) therefore continual handling of mice throughout could have impacted disease severity. Model selection is important in pre-clinical DMD studies, and we expected aggravation would induce a more human comparable phenotype. The use of aged *mdx* mice, which do show human relevant characteristics particularly regarding fibrosis and loss of function, will be crucial for well-informed clinical translation decisions.

In summary, our data highlight DMF could be a promising translational candidate for DMD with a favourable safety profile. Our follow-up study due for completion in 2025 intends to confirm longitudinal, disease modifying capacity as a final hurdle to clinical translation. Moderate-term DMF demonstrated anti-inflammatory, anti-fibrotic, and anti-lipogenic effects against the mild DMD phenotype of 8-week-old exercise aggravated *mdx* mice. A strong point of our study was the inclusion of a combinatorial DMF+PRED group to survey potential drug interactions. Worsening of skeletal muscle fibrosis with combined treatment suggests careful consideration of dose and dosing regimen will be needed during clinical trial design.

## Methods

### Animals

Male C57BL/10ScSnJ (WT) and dystrophin-deficient C57BL/10ScSn-*Dmd^mdx^/*J (*mdx*) mice were bred at Western Centre for Health, Research and Education (Sunshine Hospital, Victoria, Australia). Animals were on a 12:12 hour light-dark cycle at 20-25°C and 40% humidity. Litters were weaned at 21 days of age and randomly assigned to cages (up to 4/cage). Food and water consumption was monitored weekly (Supplemental Figure 5A-D).

### Treatment

*Mdx* mice were randomly assigned to treatment groups at 21 days of age: (1) 0.5% methylcellulose (vehicle), (2) DMF, (3) PRED or (4) a DMF and PRED combination (all treatments suspended in vehicle). WT mice were treated with 0.5% methylcellulose vehicle. Animals were weighed and treated daily up until 8 weeks of age via oral gavage. Treatments were prepared relative to individual body weights to give a final daily dosage of 100mg.kg^-1^/day DMF, 5mg.kg^-1^/day PRED or 100mg.kg^-1^/day DMF with 5mg.kg^-1^/day PRED. Selected dosages are consistent with our previous study (16), which aligns with pre-clinical studies assessing DMF for MS (67) and work by Manico *et al.* (68) who pre-clinically used this PRED dosage in *mdx* mice. There was no impact of any treatment on body weight or food and water consumption in *mdx* mice (Supplemental Figure 5A-D).

### Treadmill exercise-aggravation protocol

From 4 weeks of age, mice in the aggravated cohort were subjected to a 30 minute run on a horizontal treadmill (PanLab Harvard Bioscience) at 12 m/min, twice-weekly based on the protocol developed by TREAT-NMD (SOP DMD_M.2.1.001) (19). All animals were exercised in a temperature and light controlled environment and the exercise was performed with constant monitoring. A forced treadmill run to fatigue, starting at 5 m/min speed for 5 minutes and increasing by 1 m/min thereafter, was substituted for the final treadmill run and was undertaken 3 days prior to the experimental endpoint.

### Functional muscle strength and neuromotor coordination testing

Forelimb and whole-body grip strength was assessed weekly as previously stated (16). Neuromotor coordination was measured across 3 trials on a Rotarod (Ugo Basile, VA, Italy) at the experimental endpoint at 8 weeks of age. Mice were acclimatised one day prior to the test and each trial started with a stabilizing period followed by acceleration (up to 45rpm) until either the animal fell from the rotarod, or 600 seconds was reached (20). Time to fall (in seconds) was recorded for each mouse. All functional testing was performed blinded, by the same experimenter. Endpoint data are shown.

### Respiratory function

Respiratory function was assessed once at the experimental endpoint at 8 weeks of age in conscious, unrestrained mice using a whole-body, non-invasive plethysmograph (WBP; Data Sciences International, USA). The plethysmograph and bias flow regulator were calibrated prior to the run. Mice were placed in one of the four plethysmograph chambers and acclimated for 30 mins. Thereafter, respiratory measurements were collected using Buxco FinePointe software over a 15 min period.

### Open field animal activity testing

Open field testing enabled us to test ambulation and locomotor function at the experimental endpoint at 8 weeks of age. The test utilises a square, acrylic box (720 mm x 720 mm and 330 mm high) in a light and temperature-controlled room. At 8 weeks of age, animals were placed in the centre of the box and left to freely roam for 10 minutes. The test is captured from above and is automatically analysed via the TopScan LITE System utilising the animals centre of mass to track it over the course of the experiment.

### Body composition analysis

At 8 weeks of age, body composition was analysed using the EchoMRI-100H scanner (EchoMRI, Houston, USA) as conducted by us previously (69, 70).

### Kyphosis Index

Prior to endpoint, at 8 weeks of age, animals were lightly anaesthetised (2.5% isoflurane) and X-rayed in an IVIS Lumina X5 (PerkinElmer). Images were acquired with Living Image(R) 4.8.0 (64bit) using photograph (medium binning, F/stop 6) and X-ray (high resolution binning, F/stop 6) overlay. Images were imported into ImageJ and the kyphosis index was calculated from a line drawn between the C7 vertebra to L6 (usually corresponding with the posterior edge of the iliac wing; line AB) divided by a line perpendicular to this from the dorsal edge of the vertebra at the point of greatest curvature (line CD) (71).

### Biomarkers

#### Urine

Before functional testing at 8 weeks of age, mice were temporarily housed in sterile individual holding cages and urine was collected and snap frozen. Oxidative stress biomarker, 8-isoprostane, was quantitated via enzyme-linked immunoassay (ELISA; Cayman Chemical Company, USA).

#### Blood

Blood was collected from the endpoint surgery via cardiac puncture into Greiner Bio-One 0.5ml lithium heparin tubes. Plasma was separated via centrifugation (5 mins, 3000 x g, 4°C). Creatine kinase units were quantitated spectrophotometrically (CK-NAC kit, Randox Laboratories, UK) (72).

### Surgical Procedure

At the experimental endpoint, animals were weighed, treated via gavage, and then deeply anaesthetised (4% induction and 2.5% maintenance isoflurane). FDBs were removed, the left EDL and soleus were tied tendon to tendon with suture thread and excised for *ex vivo* contractile experimentation. Other muscles and organs were excised and snap frozen to be weighed later (refer to Supplemental Tables 1 and 2).

### *Ex vivo* Contractile Function

After excision, the proximal ends of the EDL and soleus muscles were placed onto a force transducer and the distal end fixed to a micromanipulator in organ baths (Danish Myo Technology, Hinnerup, Denmark) via knotted loops. In depth information regarding the contractile protocol has previously been described by our lab group (73). Outcome measures included specific/absolute force, force frequency, fatigue, and recovery.

### Seahorse Extracellular Flux

Excised FDB’s were prepared and incubated as previously described by us (74) using a standard mitochondrial stress test on a Seahorse Bioscience XF24 Analyser (Agilent).

### Histological analyses of the quadriceps and heart

The right quadriceps and hearts were initially snap frozen then later slow thawed at 4°C in 10% neutral buffered formalin for 48 hours. Following fixation, quadriceps and hearts were transferred to 70% ethanol until paraffin embedding. Sections were cut at 5 µm thick for the quadriceps and heart. Hearts were cut diagonally at the mid-ventricle. Sections were stained and assessed as detailed below. All assessments were performed on blinded images by the same experimenter. Full cross-sectional areas were imaged at 1X using a Zeiss Axio Imager Z2 microscope (Carl Zeiss MicroImaging, GmbH, Germany).

#### Haematoxylin and Eosin (H&E)

Quadriceps and hearts were stained with H&E (74). For quantification of healthy versus unhealthy tissue in the quadriceps, the H&E colour deconvolution plugin on ImageJ was used. The eosin component (normal, undamaged myofibers) was measured via thresholding (TREAT-NMD SOP: DMD_M.1.2.007) (75). The proportion of centronucleated fibres in the quadriceps was calculated by analysing ~200 fibres per section (76, 77). For active cardiomyocyte degeneration of the heart, degenerating tissue and clusters of inflammatory infiltrate were measured and expressed relative to the total area (78).

#### Masson’s Trichrome

Masson’s trichrome was used to assess collagen/fibrotic connective tissue content in the quadriceps and heart. Analysis of the percent area of fibrosis in both the quadriceps and heart was conducted using the Masson’s colour deconvolution plugin on ImageJ (33).

#### Immunohistochemistry

To quantify macrophage and adipocyte infiltration, anti-CD68 (ab125212) and perilipin-1 (9349S) primary antibodies were utilised at a dilution of 1:500 and 1:200, respectively and stained as described by us previously (16). CD68 positive macrophages were manually counted using DotDotGoose (version 1.7.0) and expressed as number per square millimetre of muscle cross section as measured via ImageJ. Adipocytes were manually traced using ImageJ and expressed as percentage of the total area.

### Transcriptomics

#### RNA extractions, quantification, and integrity

Frozen quadriceps muscles (10 to 15 mg) (*n*=10 per group) were removed from −80C° storage where extractions were performed using the AllPrep DNA/RNA/miRNA Universal Kit (Qiagen, Valencia, USA) which includes a genomic DNA elimination step to remove DNA from total RNA. Aliquots were taken for RNA quantification and integrity assessment using a Nanodrop spectrophotometer (Thermo Fisher Scientific). 1 μl of sample was placed on the instrument and readings for RNA concentration (ng/μl), the A260/280 ratio, and A260/230 ratio were recorded. Samples were diluted to 100 +/- 20ng/μl and RNA integrity was assessed using a TapeStation System 4150 (Agilent Technologies, USA). An RNA quality indicator score of greater than 7 was considered intact and of high integrity (79).

#### RNA sequencing

The library construction and RNA sequencing were performed by Beijing Genomics Institute (Shenzhen, China) and ~1μg total RNA was used for library construction. For library preparation, samples were denatured, and mRNA was enriched using oligo (dT) attached magnetic beads. The mRNA was fragmented into small pieces and the double strand cDNA was synthesised. These double-stranded cDNA fragments were subjected to end-repair and a single ‘A’ nucleotide is added to the 3’ ends of the blunt fragments and subsequently ligated to the adapter. The ligation product was purified and enriched with PCR amplification to yield the final cDNA library. Sufficient quality DNA nanoballs (DNBs) were loaded into patterned nanoarrays using a high-intensity DNA nanochip technique and sequenced through combinatorial Probe-Anchor Synthesis (cPAS).

#### Bioinformatics

FASTQ files were processed using the NfCore/RNAseq (v3.10.1) pipeline (80). Reads were aligned to the *Mus musculus* Ensembl GRCm39.release-109 reference genome using STAR aligner (81) and quantified using Salmon (82) producing raw genes count matrix. Various quality control metrics were generated and summarised in a MultiQC report (83). Raw counts were then analysed with Degust (84), a web tool that performs normalisation using trimmed mean of M-values (TMM) (85), and differential expression analysis performed using *limma* (86) and *voom* (87).

Aggravated *mdx* model was expressed relative to WT exercised control to rule out the effects of exercise-related adaptations. The data presented in heatmaps was considered statistically significantly different based on a threshold of a log^-2^ fold-change (log_2_FC) of ±>1.0 (2-fold difference with a corresponding FDR of <0.05).

### Metabolomics and Lipidomics

Metabolomics was run and analysed at Metabolomics Australia (Bio21 Molecular Science and Biotechnology Institute, University of Melbourne, Parkville, Australia).

#### Extraction and sample preparation

Tissue optimisation was performed and 30mg of tissue was determined to be optimal biomass of tissue required to provide the best metabolite coverage in samples. Tissues were pre-weighed and extracted by Handheld homogenisation (Astral Scientific, Australia) with pulsing at 13,000 rpm (6 x 10 seconds) in 300 μl of ice-cold 3:1 Methanol:Milli-Q water containing MA-Internal Standards. Extracts were vortexed for 10 seconds, mixed on a thermomixer and centrifuged at 4°C for 10 minutes at maximum speed.

#### Polar metabolite analysis

Analyses of the polar analytes in the samples were performed on the Orbitrap ID-X Tribrid mass spectrometer (Thermo Scientific, Waltham, MA, USA) coupled to a Vanquish Horizon UHPLC system (Thermo Scientific, Waltham, MA, USA). 100 μl of the supernatant was pipetted into a glass HPLC vial containing an insert. A further 20 μl from each sample was pooled to generate a pooled biological quality control sample (pbQC), which was run after every five-biological samples.

#### Targeted lipid analysis

The remaining homogenate was further extracted in 300 μl of 100% chloroform (containing MA-lipid Internal Standards) was added to create a ratio of 2:1 chloroform:methanol for the remaining homogenate. Extracts were vigorously vortexed to resuspend the tissue pellet. Samples were mixed on a thermomixer and centrifuged at 10°C for 5 mins at 25,155 x g. The LCMS methodologies for polar metabolite and targeted lipid analyses are outlined previously (88).

#### Bioinformatics

Statistical data analysis and data normalisation of metabolomics and lipidomics was performed in MetaboAnalyst 6.0. Data were normalised by median log transformation without data scaling. 2.0-fold change threshold and a raw *P* value of <0.05 considered significant for data presented in heatmaps.

### Statistics

Data are presented as mean ± SEM unless otherwise stated. One-way ANOVAs were performed using Prism software (GraphPad, La Jolla, CA) for all analyses unless otherwise stated to assess either the extent of phenotype or *mdx* VEH relative to treatment groups and Tukey’s post hoc test was used for multiple comparisons. For mitochondrial oxygen consumption/extracellular flux and contractile force-frequency experiments, a two-way ANOVA with repeated measures was performed using Sidak posthoc testing to interrogate differences between phenotype/treatment and time/frequency. A *P* < 0.05 was considered significant and trends were reported at *P* < 0.1. Outliers were removed if outside ± 2 standard deviations from the mean.

### Study Approval

Ethics approval was granted by Victoria University Animal Ethics Committee. All breeding, handling, and animal experiments adhered to the Australian Code of Practice for Care and Use of Animals for Scientific Purposes (National Health and Medical Research Council, Australia, 8^th^ edition). At 14 days of age, mice were transferred from breeding program (AEC# 20-005) to an approved experimental project (AEC# 20-006).

## Data availability

All individual data values represented in graphs and are available in the supporting data values file.

## Author contributions

CAT, DF, and ER conceptualised and designed the study. SK, CAT, RMB, BQ, BAA, and ER conducted animal studies. SK, CAT, RMB, GS, DD, and ER conducted experiments. SK, CAT, RMB, RB, GS, NK, ALP, TJY, and ER conducted data analysis. CAT, NK, AP, XY, NS, JH, BN, MVP, AAR, DDL, DF and ER contributed to the methodological and technical know-how, resources, and data interpretation. SK, CAT, and ER drafted the manuscript. All authors reviewed and approved the final manuscript.

## Acknowledgments

This work was supported by funding from The Muscular Dystrophy Association U.S.A (Ideas grant: MDA871929) and Australian Physiological Society (AuPS; HDR Student Small Grant). ER’s lab is supported by AFM Téléthon (France), The Jack Brockhoff Foundation and Duchenne Parent Project Netherlands. SK, CAT, and ER acknowledge outstanding support from Anne Luxford, Tricia Murphy, and Steve Holloway with animal care and breeding (Victoria University Animal Services, WCHRE, Sunshine Hospital). We gratefully acknowledge the Melbourne Histology Platform, The University of Melbourne, for expert service in tissue embedding, cutting, and staining. We gratefully acknowledge Patricia Quiambao from Metabolomics Australia, University of Melbourne, for assistance with metabolomics studies. We gratefully acknowledge Annemieke Aartsma-Rus and Maaike Van Putten, Leiden University Medical Centre, for their advice on the exercise aggravation protocol and critical reading of the manuscript. Schematics were created with BioRender.com and all figures were constructed in Adobe Illustrator.

## References

1. Duan D, Goemans N, Takeda Si, Mercuri E, and Aartsma-Rus A. Duchenne muscular dystrophy. Nature Reviews Disease Primers. 2021;7(1):13.

2. Wong SH, McClaren BJ, Archibald AD, Weeks A, Langmaid T, Ryan MM, et al. A mixed methods study of age at diagnosis and diagnostic odyssey for Duchenne muscular dystrophy. Eur J Hum Genet. 2015;23(10):1294–300.

3. Wahlgren L, Kroksmark A-K, Lindblad A, Tulinius M, and Sofou K. Respiratory comorbidities and treatments in Duchenne muscular dystrophy: impact on life expectancy and causes of death. Journal of Neurology. 2024.

4. Mhandire DZ, Burns DP, Roger AL, O’Halloran KD, and ElMallah MK. Breathing in Duchenne muscular dystrophy: translation to therapy. J Physiol. 2022;600(15):3465–82.

5. Passamano L, Taglia A, Palladino A, Viggiano E, D’Ambrosio P, Scutifero M, et al. Improvement of survival in Duchenne Muscular Dystrophy: retrospective analysis of 835 patients. Acta Myol. 2012;31(2):121–5.

6. Archer JE, Gardner AC, Roper HP, Chikermane AA, and Tatman AJ. Duchenne muscular dystrophy: the management of scoliosis. J Spine Surg. 2016;2(3):185–94.

7. Szwec S, Kapłucha Z, Chamberlain JS, and Konieczny P. Dystrophin- and Utrophin-Based Therapeutic Approaches for Treatment of Duchenne Muscular Dystrophy: A Comparative Review. BioDrugs. 2024;38(1):95–119.

8. Lek A, Wong B, Keeler A, Blackwood M, Ma K, Huang S, et al. Death after High-Dose rAAV9 Gene Therapy in a Patient with Duchenne’s Muscular Dystrophy. New England Journal of Medicine. 2023;389(13):1203–10.

9. Deng J, Zhang J, Shi K, and Liu Z. Drug development progress in duchenne muscular dystrophy. Front Pharmacol. 2022;13:950651.

10. Tulangekar A, and Sztal TE. Inflammation in Duchenne Muscular Dystrophy-Exploring the Role of Neutrophils in Muscle Damage and Regeneration. Biomedicines. 2021;9(10).

11. Kourakis S, Timpani CA, Campelj DG, Hafner P, Gueven N, Fischer D, et al. Standard of care versus new-wave corticosteroids in the treatment of Duchenne muscular dystrophy: Can we do better? Orphanet Journal of Rare Diseases. 2021;16(1):117.

12. Smith EC, Conklin LS, Hoffman EP, Clemens PR, Mah JK, Finkel RS, et al. Efficacy and safety of vamorolone in Duchenne muscular dystrophy: An 18-month interim analysis of a non-randomized open-label extension study. PLoS Med. 2020;17(9):e1003222.

13. Mah JK, Clemens PR, Guglieri M, Smith EC, Finkel RS, Tulinius M, et al. Efficacy and Safety of Vamorolone in Duchenne Muscular Dystrophy: A 30-Month Nonrandomized Controlled Open-Label Extension Trial. JAMA Netw Open. 2022;5(1):e2144178.

14. Mercuri E, Vilchez JJ, Boespflug-Tanguy O, Zaidman CM, Mah JK, Goemans N, et al. Safety and efficacy of givinostat in boys with Duchenne muscular dystrophy (EPIDYS): a multicentre, randomised, double-blind, placebo-controlled, phase 3 trial. The Lancet Neurology. 2024;23(4):393–403.

15. Kourakis S, Timpani CA, de Haan JB, Gueven N, Fischer D, and Rybalka E. Targeting Nrf2 for the treatment of Duchenne Muscular Dystrophy. Redox Biol. 2021;38:101803.

16. Timpani CA, Kourakis S, Debruin DA, Campelj DG, Pompeani N, Dargahi N, et al. Dimethyl fumarate modulates the dystrophic disease program following short-term treatment. JCI Insight. 2023;8(21).

17. Willmann R, De Luca A, Benatar M, Grounds M, Dubach J, Raymackers JM, et al. Enhancing translation: guidelines for standard pre-clinical experiments in mdx mice. Neuromuscul Disord. 2012;22(1):43–9.

18. Schill KE, Altenberger AR, Lowe J, Periasamy M, Villamena FA, Rafael-Fortney JA, et al. Muscle damage, metabolism, and oxidative stress in mdx mice: Impact of aerobic running. Muscle Nerve. 2016;54(1):110–7.

19. De Luca A. Use of treadmill and wheel exercise for impact on mdx mice phenotype (SOP DMD_M.2.1.001). https://www.treat-nmd.org/wp-content/uploads/2023/07/MDX-DMD_M.2.1.001.pdf.

20. Aartsma-Rus A, and van Putten M. Assessing functional performance in the mdx mouse model. J Vis Exp. 2014 (85).

21. Lavi E, Cohen A, Libdeh AA, Tsabari R, Zangen D, and Dor T. Growth hormone therapy for children with Duchenne muscular dystrophy and glucocorticoid induced short stature. Growth Horm IGF Res. 2023;72-73:101558.

22. Ward LM, and Weber DR. Growth, pubertal development, and skeletal health in boys with Duchenne Muscular Dystrophy. Curr Opin Endocrinol Diabetes Obes. 2019;26(1):39–48.

23. Suárez-Calvet X, Fernández-Simón E, Natera D, Jou C, Pinol-Jurado P, Villalobos E, et al. Decoding the transcriptome of Duchenne muscular dystrophy to the single nuclei level reveals clinical-genetic correlations. Cell Death Dis. 2023;14(9):596.

24. Clarkson PM, Devaney JM, Gordish-Dressman H, Thompson PD, Hubal MJ, Urso M, et al. ACTN3 genotype is associated with increases in muscle strength in response to resistance training in women. Journal of Applied Physiology. 2005;99(1):154–63.

25. Pappas CT, Krieg PA, and Gregorio CC. Nebulin regulates actin filament lengths by a stabilization mechanism. Journal of Cell Biology. 2010;189(5):859–70.

26. Lombardo SD, Basile MS, Ciurleo R, Bramanti A, Arcidiacono A, Mangano K, et al. A Network Medicine Approach for Drug Repurposing in Duchenne Muscular Dystrophy. Genes (Basel). 2021;12(4).

27. Hanna S, Xianzhen H, David KC, Ying S, Matthew SA, Merry-Lynn NM, et al. Late-Stage Skeletal Muscle Transcriptome in Duchenne muscular dystrophy shows a BMP4-Induced Molecular Signature. bioRxiv. 2024:2024.04.19.590266.

28. Vianello S, Pantic B, Fusto A, Bello L, Galletta E, Borgia D, et al. SPP1 genotype and glucocorticoid treatment modify osteopontin expression in Duchenne muscular dystrophy cells. Human Molecular Genetics. 2017;26(17):3342–51.

29. Howard ZM, Gomatam CK, Rabolli CP, Lowe J, Piepho AB, Bansal SS, et al. Mineralocorticoid receptor antagonists and glucocorticoids differentially affect skeletal muscle inflammation and pathology in muscular dystrophy. JCI Insight. 2022;7(19).

30. Ramshanker N, Aagaard M, Hjortebjerg R, Voss TS, Møller N, Jørgensen JOL, et al. Effects of Prednisolone on Serum and Tissue Fluid IGF-I Receptor Activation and Post-Receptor Signaling in Humans. The Journal of Clinical Endocrinology & Metabolism. 2017;102(11):4031–40.

31. Hathout Y, Liang C, Ogundele M, Xu G, Tawalbeh SM, Dang UJ, et al. Disease-specific and glucocorticoid-responsive serum biomarkers for Duchenne Muscular Dystrophy. Sci Rep. 2019;9(1):12167.

32. Bhargava P, Fitzgerald KC, Venkata SLV, Smith MD, Kornberg MD, Mowry EM, et al. Dimethyl fumarate treatment induces lipid metabolism alterations that are linked to immunological changes. Ann Clin Transl Neurol. 2019;6(1):33–45.

33. Morroni J, Schirone L, Vecchio D, Nicoletti C, D’Ambrosio L, Valenti V, et al. Accelerating the Mdx Heart Histo-Pathology through Physical Exercise. Life. 2021;11(7):706.

34. SOGC Guideline Retirement Notice No. 2. J Obstet Gynaecol Can. 2022;44(10):1104–12.

35. Culbreth M, and Aschner M. GSK-3β, a double-edged sword in Nrf2 regulation: Implications for neurological dysfunction and disease. F1000Res. 2018;7:1043.

36. Nakae Y, Dorchies OM, Stoward PJ, Zimmermann BF, Ritter C, and Ruegg UT. Quantitative evaluation of the beneficial effects in the mdx mouse of epigallocatechin gallate, an antioxidant polyphenol from green tea. Histochemistry and Cell Biology. 2012;137(6):811–27.

37. Fiore PF, Benedetti A, Sandonà M, Madaro L, De Bardi M, Saccone V, et al. Lack of PKCθ Promotes Regenerative Ability of Muscle Stem Cells in Chronic Muscle Injury. Int J Mol Sci. 2020;21(3).

38. Grzegorzewska AP, Seta F, Han R, Czajka CA, Makino K, Stawski L, et al. Dimethyl Fumarate ameliorates pulmonary arterial hypertension and lung fibrosis by targeting multiple pathways. Scientific Reports. 2017;7(1):41605.

39. Kato K, Papageorgiou I, Shin YJ, Kleinhenz JM, Palumbo S, Hahn S, et al. Lung-Targeted Delivery of Dimethyl Fumarate Promotes the Reversal of Age-Dependent Established Lung Fibrosis. Antioxidants (Basel). 2022;11(3).

40. Liu J, Wu Z, Liu Y, Zhan Z, Yang L, Wang C, et al. ROS-responsive liposomes as an inhaled drug delivery nanoplatform for idiopathic pulmonary fibrosis treatment via Nrf2 signaling. J Nanobiotechnology. 2022;20(1):213.

41. Oh CJ, Kim J-Y, Choi Y-K, Kim H-J, Jeong J-Y, Bae K-H, et al. Dimethylfumarate Attenuates Renal Fibrosis via NF-E2-Related Factor 2-Mediated Inhibition of Transforming Growth Factor-β/Smad Signaling. PLOS ONE. 2012;7(10):e45870.

42. Laurila PP, Luan P, Wohlwend M, Zanou N, Crisol B, Imamura de Lima T, et al. Inhibition of sphingolipid de novo synthesis counteracts muscular dystrophy. Sci Adv. 2022;8(4):eabh4423.

43. Radley-Crabb HG, Marini JC, Sosa HA, Castillo LI, Grounds MD, and Fiorotto ML. Dystropathology increases energy expenditure and protein turnover in the mdx mouse model of duchenne muscular dystrophy. PLoS One. 2014;9(2):e89277.

44. Mao C, Gao M, Zang S-K, Zhu Y, Shen D-D, Chen L-N, et al. Orthosteric and allosteric modulation of human HCAR2 signaling complex. Nature Communications. 2023;14(1):7620.

45. Molina T, Fabre P, and Dumont NA. Fibro-adipogenic progenitors in skeletal muscle homeostasis, regeneration and diseases. Open Biol. 2021;11(12):210110.

46. Mandal N, Grambergs R, Mondal K, Basu SK, Tahia F, and Dagogo-Jack S. Role of ceramides in the pathogenesis of diabetes mellitus and its complications. J Diabetes Complications. 2021;35(2):107734.

47. Miranda ER, and Funai K. Suppression of de novo sphingolipid biosynthesis mitigates sarcopenia. Nat Aging. 2022;2(12):1088–9.

48. Dasgupta S, and Ray SK. Insights into abnormal sphingolipid metabolism in multiple sclerosis: targeting ceramide biosynthesis as a unique therapeutic strategy. Ther Targets Neurol Dis. 2017;4.

49. De la Garza-Rodea AS, Moore SA, Zamora-Pineda J, Hoffman EP, Mistry K, Kumar A, et al. Sphingosine Phosphate Lyase Is Upregulated in Duchenne Muscular Dystrophy, and Its Inhibition Early in Life Attenuates Inflammation and Dystrophy in Mdx Mice. Int J Mol Sci. 2022;23(14).

50. Naarding KJ, Reyngoudt H, van Zwet EW, Hooijmans MT, Tian C, Rybalsky I, et al. MRI vastus lateralis fat fraction predicts loss of ambulation in Duchenne muscular dystrophy. Neurology. 2020;94(13):e1386–e94.

51. Khattri RB, Batra A, Matheny M, Hart C, Henley-Beasley SC, Hammers D, et al. Magnetic resonance quantification of skeletal muscle lipid infiltration in a humanized mouse model of Duchenne muscular dystrophy. NMR Biomed. 2023;36(3):e4869.

52. Houwen-van Opstal SLS, Rodwell L, Bot D, Daalmeyer A, Willemsen MAAP, Niks EH, et al. BMI-z scores of boys with Duchenne muscular dystrophy already begin to increase before losing ambulation: a longitudinal exploration of BMI, corticosteroids and caloric intake. Neuromuscular Disorders. 2022;32(3):236–44.

53. Billich N, Adams J, Carroll K, Truby H, Evans M, Ryan MM, et al. The Relationship between Obesity and Clinical Outcomes in Young People with Duchenne Muscular Dystrophy. Nutrients. 2022;14(16).

54. Srivastava NK, Yadav R, Mukherjee S, and Sinha N. Perturbation of muscle metabolism in patients with muscular dystrophy in early or acute phase of disease: In vitro, high resolution NMR spectroscopy based analysis. Clin Chim Acta. 2018;478:171–81.

55. Tsonaka R, Signorelli M, Sabir E, Seyer A, Hettne K, Aartsma-Rus A, et al. Longitudinal metabolomic analysis of plasma enables modeling disease progression in Duchenne muscular dystrophy mouse models. Hum Mol Genet. 2020;29(5):745–55.

56. Dabaj I, Ferey J, Marguet F, Gilard V, Basset C, Bahri Y, et al. Muscle metabolic remodelling patterns in Duchenne muscular dystrophy revealed by ultra-high-resolution mass spectrometry imaging. Sci Rep. 2021;11(1):1906.

57. Rybalka E, Kourakis S, Bonsett CA, Moghadaszadeh B, Beggs AH, and Timpani CA. Adenylosuccinic Acid: An Orphan Drug with Untapped Potential. Pharmaceuticals. 2023;16(6):822.

58. Timpani CA, Hayes A, and Rybalka E. Revisiting the dystrophin-ATP connection: How half a century of research still implicates mitochondrial dysfunction in Duchenne Muscular Dystrophy aetiology. Med Hypotheses. 2015;85(6):1021–33.

59. Bakay M, Zhao P, Chen J, and Hoffman EP. A web-accessible complete transcriptome of normal human and DMD muscle. Neuromuscul Disord. 2002;12 Suppl 1:S125–41.

60. Zhang H, Tsui CK, Garcia G, Joe LK, Wu H, Maruichi A, et al. The extracellular matrix integrates mitochondrial homeostasis. Cell. 2024.

61. Muire PJ, Mangum LH, and Wenke JC. Time Course of Immune Response and Immunomodulation During Normal and Delayed Healing of Musculoskeletal Wounds. Front Immunol. 2020;11:1056.

62. Gloss D, Moxley RT, 3rd, Ashwal S, and Oskoui M. Practice guideline update summary: Corticosteroid treatment of Duchenne muscular dystrophy: Report of the Guideline Development Subcommittee of the American Academy of Neurology. Neurology. 2016;86(5):465–72.

63. Administration USFaD. In: Services USDoHaH ed. Rockville, MD 2005.

64. Wintzinger M, Miz K, York A, Demonbreun AR, Molkentin JD, McNally EM, et al. Effects of Glucocorticoids in Murine Models of Duchenne and Limb-Girdle Muscular Dystrophy. Methods Mol Biol. 2023;2587:467–78.

65. Quattrocelli M, Barefield DY, Warner JL, Vo AH, Hadhazy M, Earley JU, et al. Intermittent glucocorticoid steroid dosing enhances muscle repair without eliciting muscle atrophy. J Clin Invest. 2017;127(6):2418–32.

66. Lindsay A, Trewin AJ, Sadler KJ, Laird C, Della Gatta PA, and Russell AP. Sensitivity to behavioral stress impacts disease pathogenesis in dystrophin-deficient mice. The FASEB Journal. 2021;35(12):e22034.

67. Kourakis S, Timpani CA, de Haan JB, Gueven N, Fischer D, and Rybalka E. Dimethyl Fumarate and Its Esters: A Drug with Broad Clinical Utility? Pharmaceuticals (Basel). 2020;13(10).

68. Mâncio RD, Hermes TA, Macedo AB, Mizobuti DS, Rupcic IF, and Minatel E. Dystrophic phenotype improvement in the diaphragm muscle of mdx mice by diacerhein. PLoS One. 2017;12(8):e0182449.

69. Campelj DG, Debruin DA, Timpani CA, Hayes A, Goodman CA, and Rybalka E. Sodium nitrate co-supplementation does not exacerbate low dose metronomic doxorubicin-induced cachexia in healthy mice. Scientific Reports. 2020;10(1):15044.

70. Sorensen JC, Petersen AC, Timpani CA, Campelj DG, Cook J, Trewin AJ, et al. BGP-15 Protects against Oxaliplatin-Induced Skeletal Myopathy and Mitochondrial Reactive Oxygen Species Production in Mice. Front Pharmacol. 2017;8:137.

71. Laws N, and Hoey A. Progression of kyphosis in mdx mice. J Appl Physiol (1985). 2004;97(5):1970–7.

72. James C, Dugan CW, Boyd C, Fournier PA, and Arthur PG. Temporal tracking of cysteine 34 oxidation of plasma albumin as a biomarker of muscle damage following a bout of eccentric exercise. Eur J Appl Physiol. 2024.

73. Debruin DA, Timpani CA, Lalunio H, Rybalka E, Goodman CA, and Hayes A. Exercise May Ameliorate the Detrimental Side Effects of High Vitamin D Supplementation on Muscle Function in Mice. J Bone Miner Res. 2020;35(6):1092–106.

74. Timpani CA, Goodman CA, Stathis CG, White JD, Mamchaoui K, Butler-Browne G, et al. Adenylosuccinic acid therapy ameliorates murine Duchenne Muscular Dystrophy. Sci Rep. 2020;10(1):1125.

75. Grounds M. Quantification of histopathology in Haemotoxylin and Eosin stained muscle sections (SOP DMD_M.1.2.007). https://www.treat-nmd.org/wp-content/uploads/2023/07/MDX-DMD_M.1.2.007-28.pdf.

76. Stupka N, Plant DR, Schertzer JD, Emerson TM, Bassel-Duby R, Olson EN, et al. Activated calcineurin ameliorates contraction-induced injury to skeletal muscles of mdx dystrophic mice. J Physiol. 2006;575(Pt 2):645–56.

77. Stupka N, Schertzer JD, Bassel-Duby R, Olson EN, and Lynch GS. Calcineurin-Aα activation enhances the structure and function of regenerating muscles after myotoxic injury. American Journal of Physiology-Regulatory, Integrative and Comparative Physiology. 2007;293(2):R686–R94.

78. Morroni J, Schirone L, Valenti V, Zwergel C, Riera CS, Valente S, et al. Inhibition of PKCθ Improves Dystrophic Heart Phenotype and Function in a Novel Model of DMD Cardiomyopathy. International Journal of Molecular Sciences. 2022;23(4):2256.

79. Lin W, Saner NJ, Weng X, Caruana NJ, Botella J, Kuang J, et al. The Effect of Sleep Restriction, With or Without Exercise, on Skeletal Muscle Transcriptomic Profiles in Healthy Young Males. Front Endocrinol (Lausanne). 2022;13:863224.

80. Harshil Patel PE, Alexander Peltzer, Rickard Hammarén, Olga Botvinnik, Gregor Sturm, Denis Moreno, Pranathi Vemuri, silviamorins, Lorena Pantano, Friederike Hanssen, Maxime U. Garcia, Chris Cheshire, rfenouil, nf-core bot, marchoeppner, Peng Zhou, Gisela Gabernet, Daniel Straub, … Matthias Hörtenhuber.; 2021.

81. Dobin A, Davis CA, Schlesinger F, Drenkow J, Zaleski C, Jha S, et al. STAR: ultrafast universal RNA-seq aligner. Bioinformatics. 2012;29(1):15–21.

82. Patro R, Duggal G, Love MI, Irizarry RA, and Kingsford C. Salmon provides fast and bias-aware quantification of transcript expression. Nat Methods. 2017;14(4):417–9.

83. Ewels P, Magnusson M, Lundin S, and Käller M. MultiQC: summarize analysis results for multiple tools and samples in a single report. Bioinformatics. 2016;32(19):3047–8.

84. David R, Powell AP, Michael Milton. zenodo; 2019.

85. Robinson MD, and Oshlack A. A scaling normalization method for differential expression analysis of RNA-seq data. Genome Biology. 2010;11(3):R25.

86. Ritchie ME, Phipson B, Wu D, Hu Y, Law CW, Shi W, et al. limma powers differential expression analyses for RNA-sequencing and microarray studies. Nucleic Acids Research. 2015;43(7):e47-e.

87. Law CW, Chen Y, Shi W, and Smyth GK. voom: precision weights unlock linear model analysis tools for RNA-seq read counts. Genome Biology. 2014;15(2):R29.

88. Lu T, Freytag L, Narayana VK, Moore Z, Oliver SJ, Valkovic A, et al. Matrix Selection for the Visualization of Small Molecules and Lipids in Brain Tumors Using Untargeted MALDI-TOF Mass Spectrometry Imaging. Metabolites. 2023;13(11):1139.

89. Gillis JM. The recovery score to evaluate therapy efficiency in NMD: a common, quantitative and comparative scoring system (SOP DMD_M.1.1_001). https://www.treat-nmd.org/wp-content/uploads/2023/07/MDX-DMD_M.1.1_001-21.pdf.

