## Supplemental Figures and Tables for "Moderate-term dimethyl fumarate treatment reduces pathology of dystrophic skeletal and cardiac muscle in a mouse model"

**Title:** Moderate-term dimethyl fumarate treatment is anti-fibrotic in dystrophic skeletal and cardiac muscle

**Conflict of Interest:** ER discloses consultancy work for Santhera Pharmaceuticals and Epirium Bio outside of this project.

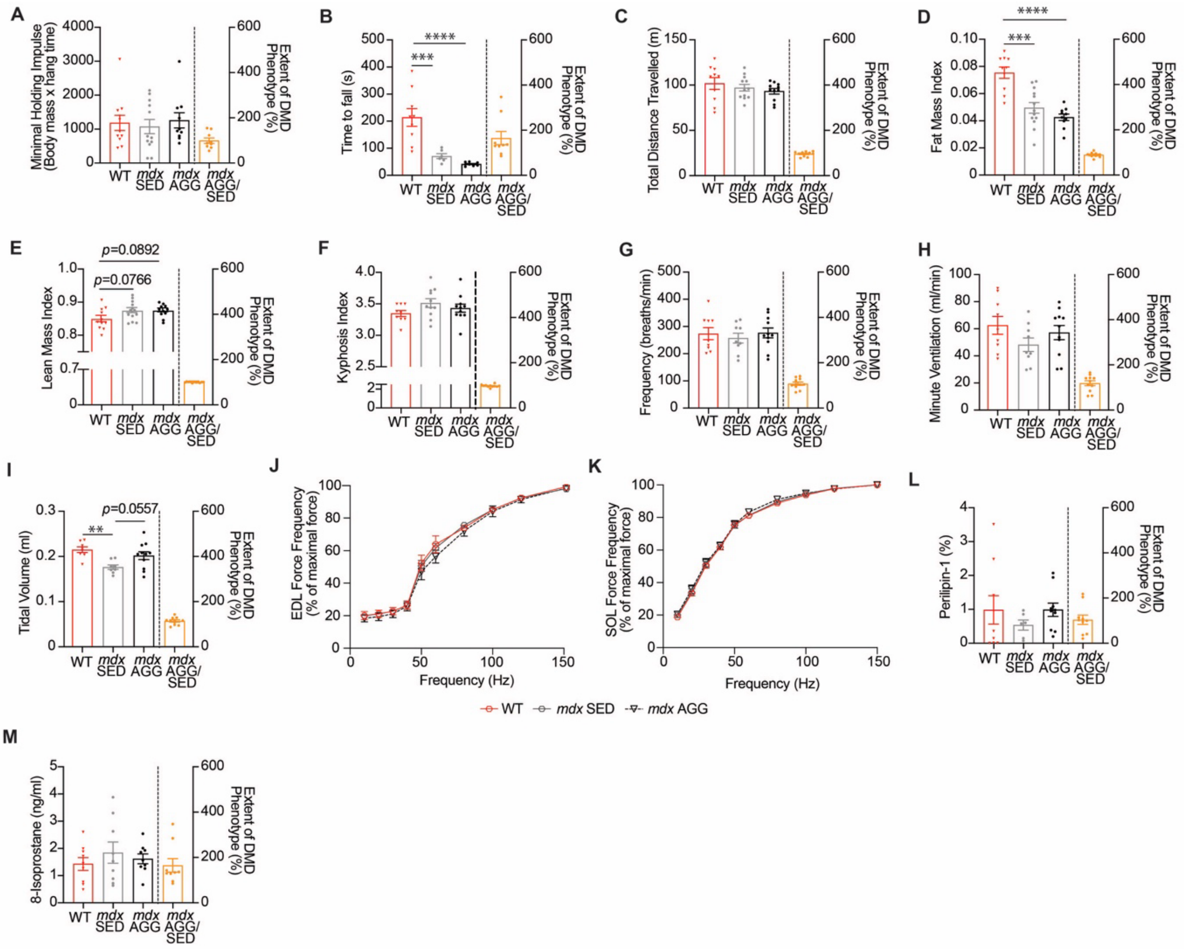


**Supplementary Figure 1. Exercise-induced aggravation has no significant effect on functional or cardiorespiratory parameters in *mdx* mice. (A)** Whole-body hang strength, **(B)** rotarod neuromotor function, **(C)** distance covered in the open field and body composition including **(D)** fat mass and **(E)** lean mass, were assessed at experimental endpoint. **(F)** The kyphosis index, a measure of thoracolumbar deformity, was calculated (17). Respiratory function indices, including **(G)** frequency, **(H)** minute ventilation and **(I)** tidal volume were measured using plethysmography. *Ex vivo* muscle contractile studies were performed and the force frequency relationship of **(J)** EDL and **(K)** SOL was determined. Percentage of **(L)** perilipin-1 positive fat deposition in the quadriceps was histologically assessed. Urinary oxidative stress biomarker was assessed via **(M)** 8-isoprostane detection. Data in **A-I**, **L-M** and are presented as mean ± SEM and *n* are indicated by individual data points. Panel **J** *n*: WT = 6, *mdx* SED = 6, *mdx* AGG = 9. Panel **K** *n*: WT = 7, *mdx* SED = 8, *mdx* AGG = 10. Statistical significance was tested via one-way ANOVA: **P*<0.05, ***P* <0.01, ****P* <0.001, *****P* <0.0001.

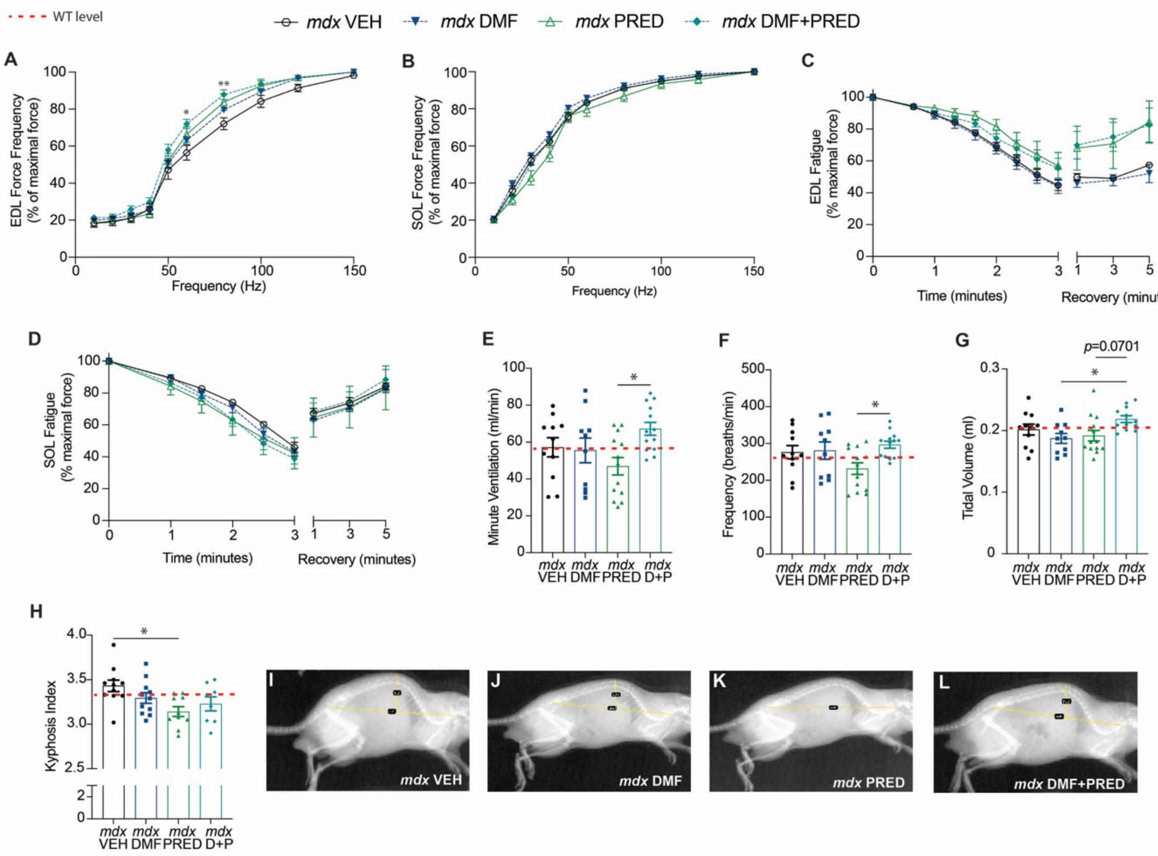


**Supplementary Figure 2. DMF has no effect on muscle contractile or respiratory properties.** The force frequency relationship in **(A)** EDL and **(B)** SOL were determined via ex *vivo* contractile experiments. The fatiguability and recovery characteristics of the **(C)** EDL and **(D)** SOL were determined. **(E)** Minute ventilation, **(F)** frequency and **(G)** tidal volume were measured via plethysmography. **(H-L)** The kyphosis index, a measure of thoracolumbar deformity, was calculated. Panel **A** *n*: *mdx* VEH = 9, *mdx* DMF =10, *mdx* PRED = 9, *mdx* DMF+PRED = 10. Panel **B** *n*: *mdx* VEH = 10, *mdx* DMF = 12, *mdx* PRED = 11, *mdx* DMF+PRED = 12. Panel **C** *n*: *mdx* VEH = 7, *mdx* DMF = 9, *mdx* PRED = 3, *mdx* DMF+PRED = 7. Panel **D** *n*: *mdx* VEH = 7, *mdx* DMF = 11, *mdx* PRED = 4, *mdx* DMF+PRED = 13. Data in **E-H** are mean ± SEM and *n* are indicated by individual data points. Statistical significance in **A-D** was tested via mixed effects analysis and in **E-H** statistical significance was tested via one-way ANOVA: *$P < 0.05$, **$P < 0.01$.

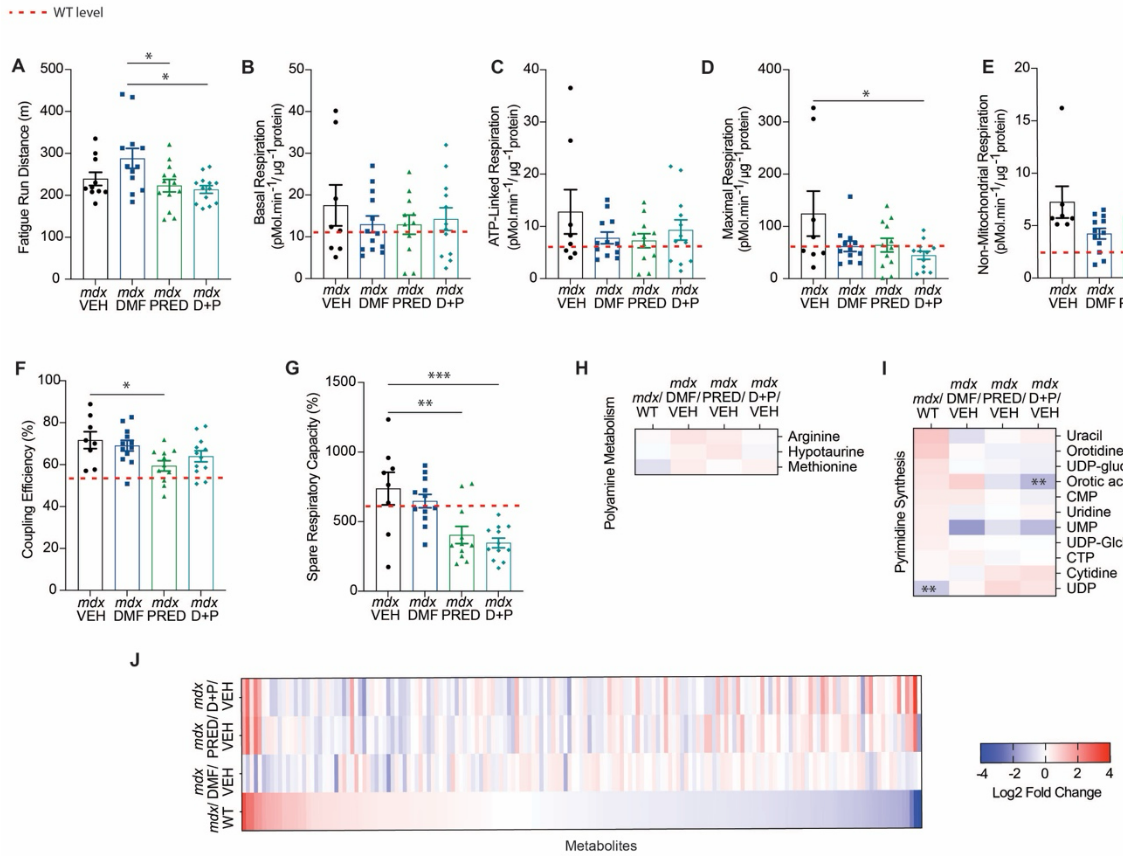


**Supplementary Figure 3. DMF has no effect on mitochondrial flexibility.** To assess mitochondrial responses to treatment, mice were subject to **(A)** a forced treadmill run to fatigue test and metabolic parameters of **(B)** basal, **(C)** ATP-linked, **(D)** maximal and **(E)** non-mitochondrial respiration were measured in isolated FDB fibres as well as the **(F)** coupling efficiency and **(G)** spare respiratory capacity. **(H)** Polyamine and **(I)** pyrimidine synthesis metabolomic pathways were probed. **(J)** Full metabolomic signature is displayed as most to least dysregulated. Data in heatmaps **(H-J)** are based on log2 fold change from WT for *mdx* VEH and *mdx* VEH for treatment groups (DMF, PRED and DMF+PRED). Data in **A-G** are presented as mean ± SEM and *n* are indicated by individual data points. Statistical significance was tested via one-way ANOVA: **P* <0.05, ***P* <0.01, ****P* <0.001.

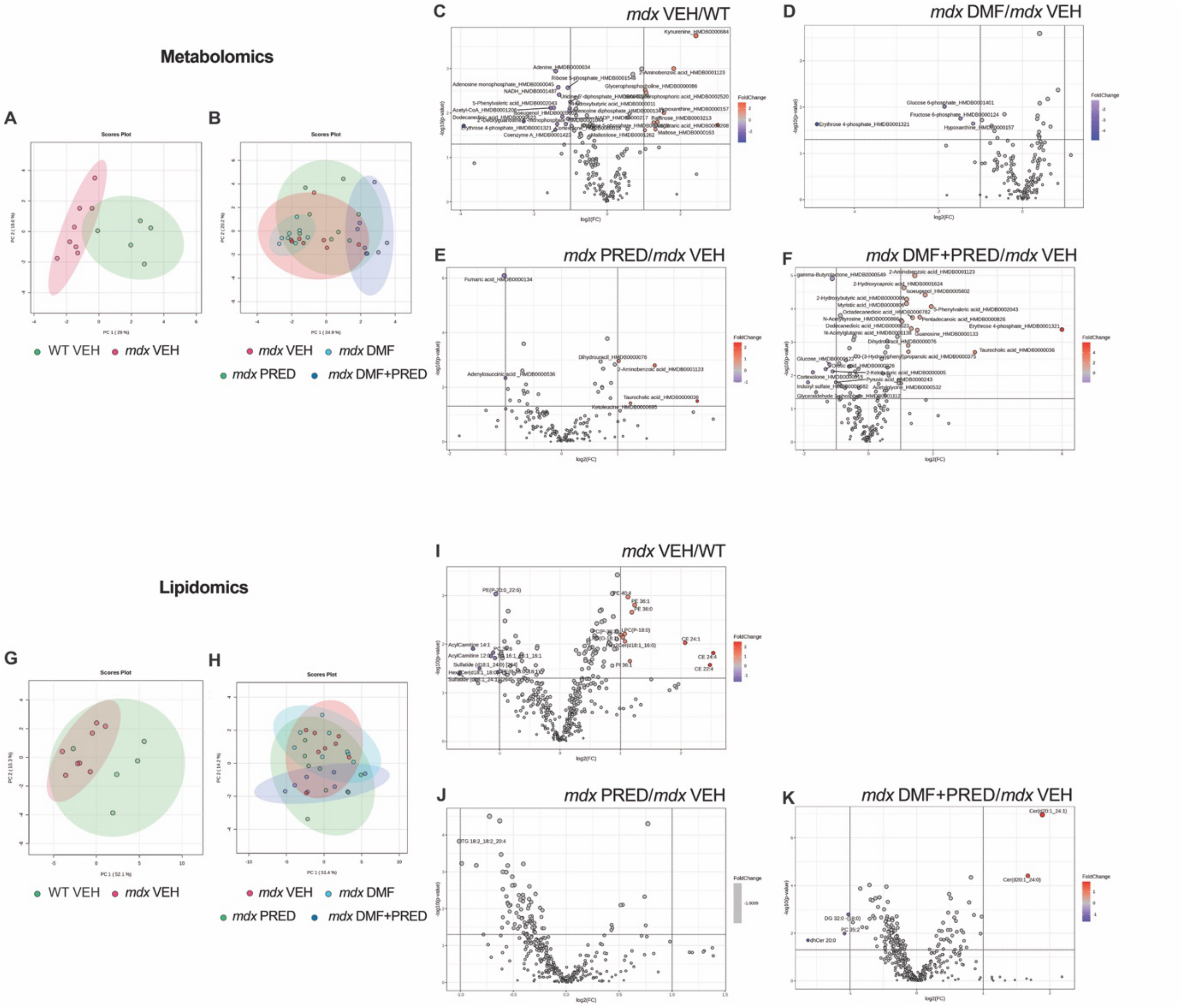


**Supplementary Figure 4. Metabolomic and lipidomic PCA and volcano plots.** PCA plots of metabolomics data showing comparisons between **(A)** WT and *mdx* VEH and **(B)** *mdx* VEH compared to treatments (DMF, PRED and DMF+PRED). Volcano plots of significant metabolomic changes in **(C)** *mdx* relative to WT **(D)** and *mdx* DMF, **(E)** PRED and **(F)** DMF+PRED relative to *mdx* VEH. PCA plots of lipidomics data showing comparisons between **(G)** WT and *mdx* VEH and **(H)** *mdx* VEH compared to treatments (DMF, PRED and DMF+PRED). Volcano plots of significant lipidomic changes in **(I)** *mdx* relative to WT **(J)** and *mdx* PRED and **(K)** DMF+PRED relative to *mdx* VEH are also shown. Note there were no significant differences in *mdx* DMF treated animals relative to *mdx* VEH.

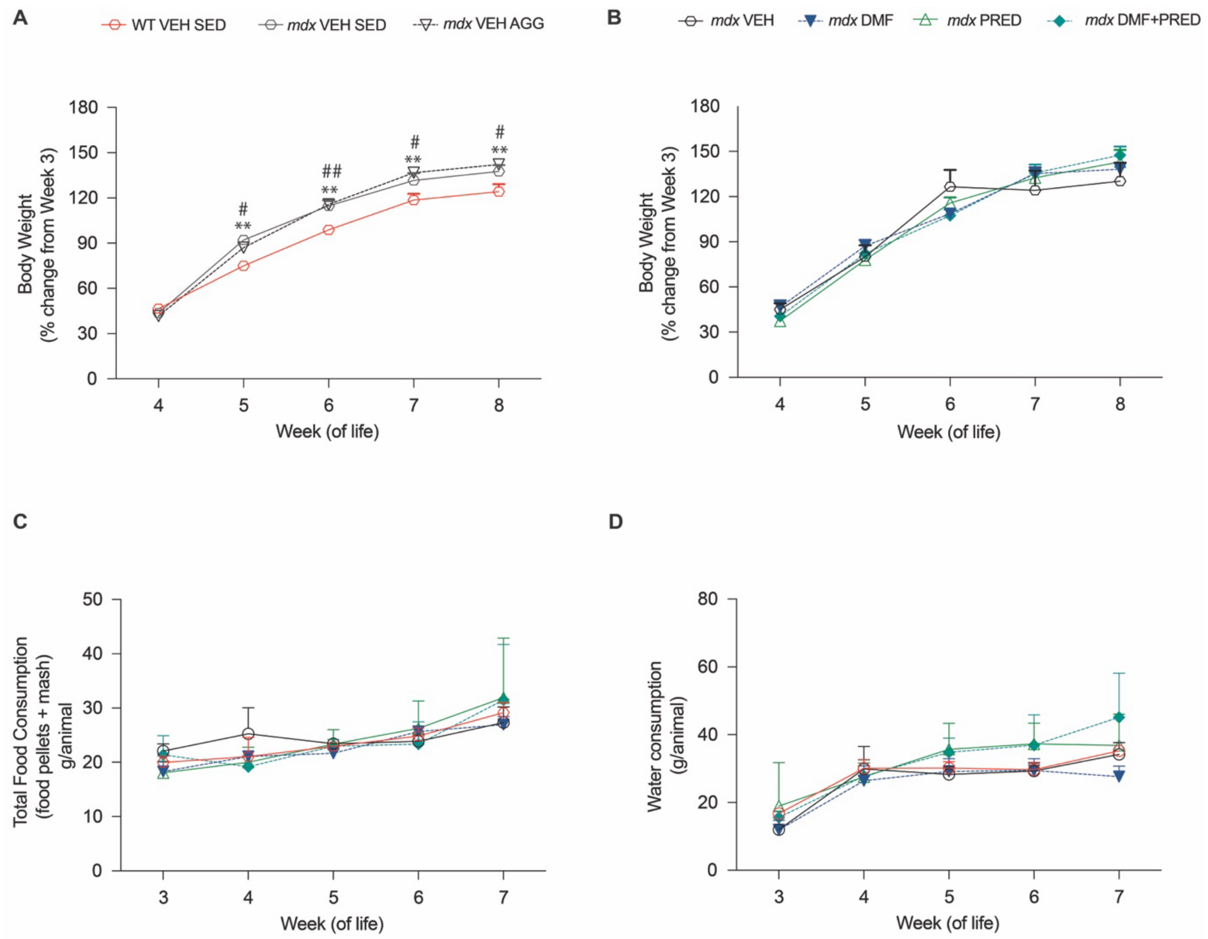


**Supplementary Figure 5. Daily DMF, PRED and DMF+PRED had no effect on body weight or food and water consumption. (A-B)** Body weight was monitored daily (presented as percentage change from beginning of experiment) whilst **(C-D)** food and water consumption was measured weekly. Panel **A** *n*: WT VEH SED = 12, *mdx* VEH SED = 12, *mdx* VEH AGG = 13. Panel **B** *n*: *mdx* VEH AGG = 13, *mdx* DMF AGG = 13, *mdx* PRED AGG = 13, *mdx* DMF+PRED AGG = 13. Panel **C** and **D** *n*: WT VEH = 23, *mdx* VEH = 19, *mdx* DMF = 23, *mdx* PRED = 21, *mdx* DMF+PRED = 28. Statistical significance was tested via two-way ANOVA: Panel A: # WT VEH vs *mdx* VEH sedentary $P<0.05$, ## WT VEH vs *mdx* VEH sedentary $P<0.01$, ** WT VEH vs *mdx* VEH aggravated $P<0.01$.

**Supplementary Table 1. Effect of phenotype and treatments on organ mass of WT VEH and dystrophic *mdx* mice.** Data are mean ± SEM. Statistical significance tested via one-way ANOVA: $^A$= *mdx* VEH significantly different from WT SED. $^B$= *mdx* VEH sedentary significantly different from *mdx* VEH aggravated. $^C$= treatment difference versus *mdx* VEH aggravated. $^D$= *mdx* DMF aggravated difference versus *mdx* PRED aggravated. $^E$= *mdx* DMF aggravated difference versus *mdx* DMF+PRED aggravated. $^F$= *mdx* PRED aggravated difference versus *mdx* DMF+PRED aggravated.

| | **WT VEH (S) (mg$^{-1}$. g bw$^{-1}$)** | ***mdx* VEH (S) (mg$^{-1}$. g bw$^{-1}$)** | ***mdx* VEH (A) (mg$^{-1}$. g bw$^{-1}$)** | ***mdx* DMF (A) (mg$^{-1}$. g bw$^{-1}$)** | ***mdx* PRED (A) (mg$^{-1}$. g bw$^{-1}$)** | ***mdx* DMF+PRED (A) (mg$^{-1}$. g bw$^{-1}$)** |
|---|---|---|---|---|---|---|
| **Heart** | 4.933 ± 0.306 (*n*=11) | 5.432 ± 0.177 (*n*=13) | 5.433 ± 0.101 (*n*=11) | 5.091 ± 0.056 (*n*=13) | 4.964 ± 0.099 (*n*=13) | 5.052 ± 0.102 (*n*=13) |
| **Lungs** | 7.560 ± 0.567 (*n*=11) | 6.483 ± 0.238 (*n*=13) | 7.111 ± 0.305 (*n*=11) | 6.778 ± 0.364 (*n*=13) | 6.913 ± 0.184 (*n*=13) | 6.34 ± 0.413 (*n*=13) |
| **Liver** | 46.245 ± 1.427 (*n*=11) | 51.918 ± 1.827$^{A, B}$ (*n*=13) | 56.030 ± 1.1765$^{A}$ (*n*=11) | 55.537 ± 1.235$^{D, E}$ (*n*=13) | 48.994 ± 0.569$^{C}$ (*n*=13) | 51.914 ± 1.067$^{C, F}$ (*n*=13) |
| **Spleen** | 3.289 ± 0.077 (*n*=11) | 3.469 ± 0.115 (*n*=13) | 3.667 ± 0.158 (*n*=11) | 3.272 ± 0.081 (*n*=13) | 2.122 ± 0.092 (*n*=13) | 2.088 ± 0.119 (*n*=13) |
| **Kidneys** | 6.801± 0.207 (*n*=11) | 7.086 ± 0.232 (*n*=13) | 7.219 ± 0.169 (*n*=11) | 7.62 ± 0.192 (*n*=13) | 6.676 ± 0.132 (*n*=13) | 7.15 ± 0.161 (*n*=13) |

**Supplementary Table 2. Effect of phenotype and treatments on muscle mass of WT VEH) and dystrophic *mdx* mice.** Data are mean ± SEM. Statistical significance tested via one-way ANOVA: $^{A}$= *mdx* VEH significantly different from WT SED. $^{B}$= *mdx* VEH sedentary significantly different from *mdx* VEH aggravated. $^{C}$= treatment difference versus *mdx* VEH aggravates. $^{D}$= *mdx* DMF aggravated difference versus *mdx* PRED aggravated. $^{E}$= *mdx* DMF aggravated difference versus *mdx* DMF+PRED aggravated. $^{F}$= *mdx* PRED aggravated difference versus *mdx* DMF+PRED aggravated.

| | **WT VEH (S) ($mg^{-1}$. g $bw^{-1}$)** | ***mdx* VEH (S) ($mg^{-1}$. g $bw^{-1}$)** | ***mdx* VEH (A) ($mg^{-1}$. g $bw^{-1}$)** | ***mdx* DMF (A) ($mg^{-1}$. g $bw^{-1}$)** | ***mdx* PRED (A) ($mg^{-1}$. g $bw^{-1}$)** | ***mdx* DMF+PRED (A) ($mg^{-1}$. g $bw^{-1}$)** |
|---|---|---|---|---|---|---|
| **Extensor digitorum longus** | 0.452 ± 0.024 (*n*=11) | 0.519 ± 0.053 (*n*=13) | 0.513 ± 0.019 (*n*=11) | 0.509 ± 0.016 (*n*=13) | 0.483 ± 0.228 (*n*=13) | 0.462 ± 0.0166 (*n*=13) |
| **Soleus** | 0.447 ± 0.053 (*n*=11) | 0.494 ± 0.038 (*n*=13) | 0.448 ± 0.016 (*n*=11) | 0.458 ± 0.0319 (*n*=13) | 0.445 ± 0.038 (*n*=13) | 0.376 ± 0.017 (*n*=13) |
| **Quadriceps** | 6.069 ± 0.269 (*n*=11) | 9.056 ± 0.601$^{A}$ (*n*=13) | 8.475 ± 0.555$^{A}$ (*n*=11) | 7.176 ± 0.280$^{C}$ (*n*=13) | 6.808 ± 0.252$^{C}$ (*n*=13) | 7.117 ± 0.207$^{C}$ (*n*=13) |
| **Gastrocnemius** | 5.515 ± 0.202 (*n*=11) | 6.759 ± 0.244$^{A}$ (*n*=13) | 6.517 ± 0.125$^{A}$ (*n*=11) | 5.936 ± 0.108 (*n*=13) | 6.099 ± 0.069 (*n*=13) | 5.839 ± 0.095$^{C}$ (*n*=13) |
| **Tibialis anterior** | 1.760 ± 0.182 (*n*=11) | 2.548 ± 0.095$^{A}$ (*n*=13) | 2.489 ± 0.050 (*n*=11) | 2.282 ± 0.046 (*n*=13) | 2.320 ± 0.063 (*n*=13) | 2.407 ± 0.051 (*n*=13) |
| **Plantaris** | 0.832 ± 0.029 (*n*=11) | 0.874 ± 0.067 (*n*=13) | 0.869 ± 0.026 (*n*=11) | 0.805 ± 0.022 (*n*=13) | 0.817 ± 0.036 (*n*=13) | 0.846 ± 0.025 (*n*=13) |
